# Beyond Simply Spinning: Improving ^1^H Resolution at Fast Magic-Angle-Spinning Frequencies Using Combined Rotation and Multiple Pulse Spectroscopy

**DOI:** 10.64898/2026.06.18.731565

**Authors:** Mrudula M. Nikam, Pragyan P. Parida, Sreejith Raran-Kurussi, P. K. Madhu, Kaustubh R. Mote

**Affiliations:** Tata Institute of Fundamental Research Hyderabad, 36/P Gopanpally Village, Serilingampally Mandal, Rangareddy District, Hyderabad-500046, Telangana, India

## Abstract

Rapid developments in magic-angle-spinning (MAS) hardware over the past two decades have made possible the acquisition of high-resolution spectra of protons in solids, fuelling studies of small and large molecules alike. Nevertheless, proton resolution, limited by the strong dipole-dipole coupling network, remains a bottleneck even at MAS frequencies exceeding 100 kHz. We present here techniques based on phase-modulated homonuclear decoupling that dramatically improve proton coherence times and resolution compared to 60–95 kHz MAS alone using low average radio-frequency amplitudes (*<* 100 kHz). A relatively high sensitivity (40– 70%) and a straightforward optimization procedure directly on the sample being studied allows these gains to be realised in large biomolecules, as demonstrated here on a 326-residue cytoskeletal protein in its filamentous state. These techniques enable experiments with improved resolution on biomolecules while simultaneously taking advantage of the higher sensitivity available on probes with relatively large rotor volumes that cannot reach higher MAS frequencies.

## II. INTRODUCTION

A high gyromagnetic ratio and a nearly 100% natural abundance makes proton an ideal nucleus for detection in nuclear magnetic resonance (NMR) experiments. Proton detection is the default in solution NMR, and is increasingly becoming common in the solid-state as well. The revolution in fast MAS probe technologies that enabled proton-detection over the past two decades has fuelled studies on a number of biomolecular systems that would have been difficult to study using traditional ^13^C detected methods [1–4]. However, the same properties that make proton an ideal nucleus for detection also lead to short coherence times and consequently, broad resonance lines in the solid state [5, 6]. The strong homonuclear dipole-dipole coupling network between protons is the primary cause of this loss in resolution. This interaction is homogeneous (as defined by Maricq and Waugh [7]), and its strength decreases only slowly with increasing MAS frequencies [5–9]. Linewidths are limited by the effect of these couplings even at spinning frequencies exceeding 100 kHz [10, 11]. It has been estimated that spinning frequencies *>*250 kHz will be required to reduce the strength of the coupling and achieve resolution comparable to that attained in perdeuterated (and partially backexchanged) proteins at the spinning frequency of 100 kHz [5]. Currently, we are more than half the way to this stage, with MAS frequencies of 160 kHz being commercially available [12, 13], and even higher spinning frequencies being reported in research labs [14].

Nevertheless, given the technical challenges in increasing MAS frequencies and the ensuing loss of sensitivity as one moves to smaller rotors to achieve this [11], several alternatives to improve resolution at a fixed MAS frequency have been explored [15–18]. Utilizing the additive averaging effect of pulsed homonuclear decoupling together with sample spinning is the oldest of these approaches, and is often called ‘Combined Rotation and Multiple Pulse Spectroscopy’ (CRAMPS). Pulsed homonuclear decoupling originates from work of Lee and Goldburg [19] and its combination with MAS was first theorized by Haeberlen and Waugh [20]. It has been implemented under MAS with several classes of pulse sequences, such as Phase-Modulated Lee-Goldburg (PMLG), Tilted Magic-Echo Sandwich (TIMES), and Decoupling Using Mind-Boggling Optimization (DUMBO) [21–25]. These have been extensively used with slow-moderate MAS frequencies (10– 30 kHz) to improve ^1^H resolution [26–29]. At MAS frequencies of 40–80 kHz, these methods need to use carefully optimized conditions to achieve the best resolution [23, 30–33]. However, these techniques are not routinely used to study biological samples at fast MAS frequencies due to the following reasons: (i) Radio-frequency (rf) pulses used in NMR experiments are far from ideal, and show a broad, non-uniform distribution in probes used for MAS experiments [34]. All implementations of CRAMPS show degraded performance in the presence of this rf-inhomogeneity. This causes an additional broadening of resonances, which is estimated to be at least 30 Hz [35]. Since linewidths in proteins approach 100– 150 Hz at MAS frequencies of 100 kHz [11], it remains unclear if the use of CRAMPS will result in net line narrowing, or if any improvement due to these sequences will be negated by the above effect. (ii) Detecting ^1^H in the direct dimension with CRAMPS requires windowed-detection, leading to an unavoidable loss in sensitivity as only a part of the signal is detected [26]. This results in an overall sensitivity *<*20%, partially negating the gains due to any improvement in resolution [26]. Even with significant line narrowing, this loss in sensitivity is unacceptably high when working with biological samples.

(iii) Aggravating the above issues, optimization procedures for CRAMPS, especially at fast MAS frequencies, are cumbersome. These are typically done on standard samples such as glycine by monitoring the directly detected spectrum and optimizing several parameters to achieve the best resolution [31–33, 36]. Such procedures do not translate well to proteins due to the large number of overlapping lines which cannot be monitored for improvement in resolution directly in a one-dimensional experiment. As such, even though applications on small molecules have been demonstrated, their translation to larger proteins has not been described.

Here, we show that all the above problems can be overcome for multidimensional experiments on biomolecules, and a resolution that is substantially better than MAS alone at the spinning frequencies of 60–95 kHz can be obtained with a relatively high sensitivity. We demonstrate this on non-deuterated and uniformly ^13^C, ^15^N labeled samples of the tripeptide formyl-methionyl-leucyl-phenylalanine (f-MLF), a microcrystalline preparation of the protein GB1 (∼ 6 kDa), and the filamentous state of a prokaryotic cytoskeletal protein ParM (∼ 36 kDa) [37– 39]. This is made possible by a technique that allows good decoupling conditions requiring low rf amplitudes to be explored systematically, and a straightforward optimization procedure directly on the sample being studied. In the following sections, we first discuss these optimization procedures and then the improvement in resolution obtained using these methods.

## III. RESULTS AND DISCUSSION

Optimizing homonuclear decoupling at spinning frequencies *>*60 kHz is complicated by the need to avoid dense resonance conditions, given by Equation 1 [30]. These correspond to zero- and first-order recoupling conditions in a Floquet theory-based analysis which takes into account two time-dependent Hamiltonians with the cycle frequencies of *v*_*r*_ (MAS frequency) and *v*_*c*_ (frequency of the homonuclear decoupling cycle) [30, 40].

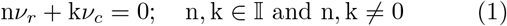

Experiments between *v*_*r*_ ∼60–80 kHz have previously shown that an optimal set up of homonuclear decoupling requires a careful optimization of the cycle time of the decoupling sequence and the rf amplitude independently [31–33]. These studies optimized the decoupling condition by monitoring ^1^H resonances using windowed acquisition. These optimizations are potentially confounded two factors: (a) inhomogeneous broadening that arises due to static and MAS-modulated rf-inhomogeneities, which depend on the rf amplitude [28, 35], and (b) the efficiency of windowed-detection, which depends on the cycle time of the decoupling sequence, a variable for optimization [26]. Additionally, it is unclear if they can be translated to larger biomolecules, where one cannot judge the quality of decoupling in a 1D experiment. Instead, we optimized homonuclear decoupling using a spin-echo following a ^15^N-filtered ^1^H experiment based on cross-polarization (Figure 1A) [41]. This set up does not suffer from the above drawbacks as spin-echos refocus inhomogeneous interactions and result in signal intensities that are dependent solely on the efficiency of the homonuclear decoupling sequence. It leads to clearly identifiable resonance conditions as well as good decoupling conditions (Figure 1, Figures S1 to S4). Homonuclear decoupling leads to the reduction in the strength of the chemical-shift interaction, described by a scaling factor (between 0 and 1). A high scaling factor is necessary to retain the resolution gain in presence of these sequences. The above approach to optimization has a singular drawback of not having direct access to this scaling factor. However, this problem is partially circumvented by the realization that the good decoupling conditions determined in this regime have a scaling factor *>*0.8 (*vide infra*). These optimizations ignore the requirement of satisfying the Lee-Goldburg condition [21, 42], as large deviations from these are anticipated at fast MAS frequencies [24, 31–33]. The range of optimization of *v*_1_ and *v*_*c*_ was chosen such that experiment does not use either very short pulses (for ensuring the stability of the sequence) or unacceptably high rf amplitudes, which may lead to deleterious heating in biological samples. Low rf amplitudes were also deemed necessary to limit inhomogeneous broadening while detection and retain any gains in resolution due to homonuclear decoupling [28]. The phase ramp consists of 3 steps, with two ramps in opposite directions making up the homonuclear decoupling shape [15]. Sequence with a higher number of steps require implementations with short pulse-lengths over which the amplitude of the pulse must stabilize, and were hence not used at these MAS frequencies. The phase modulated sequence was itself implemented as a two-step supercycle [22]. This is a critical requirement for obtaining artefact-free spectra with uniform scaling factors under MAS [22, 43]. The lower scaling factor for these sequences as compared to non-supercycled ones was not considered problematic since experimentally measured scaling factors approached a relatively high value of 0.9 (*vide infra*).

**FIG. 1.**
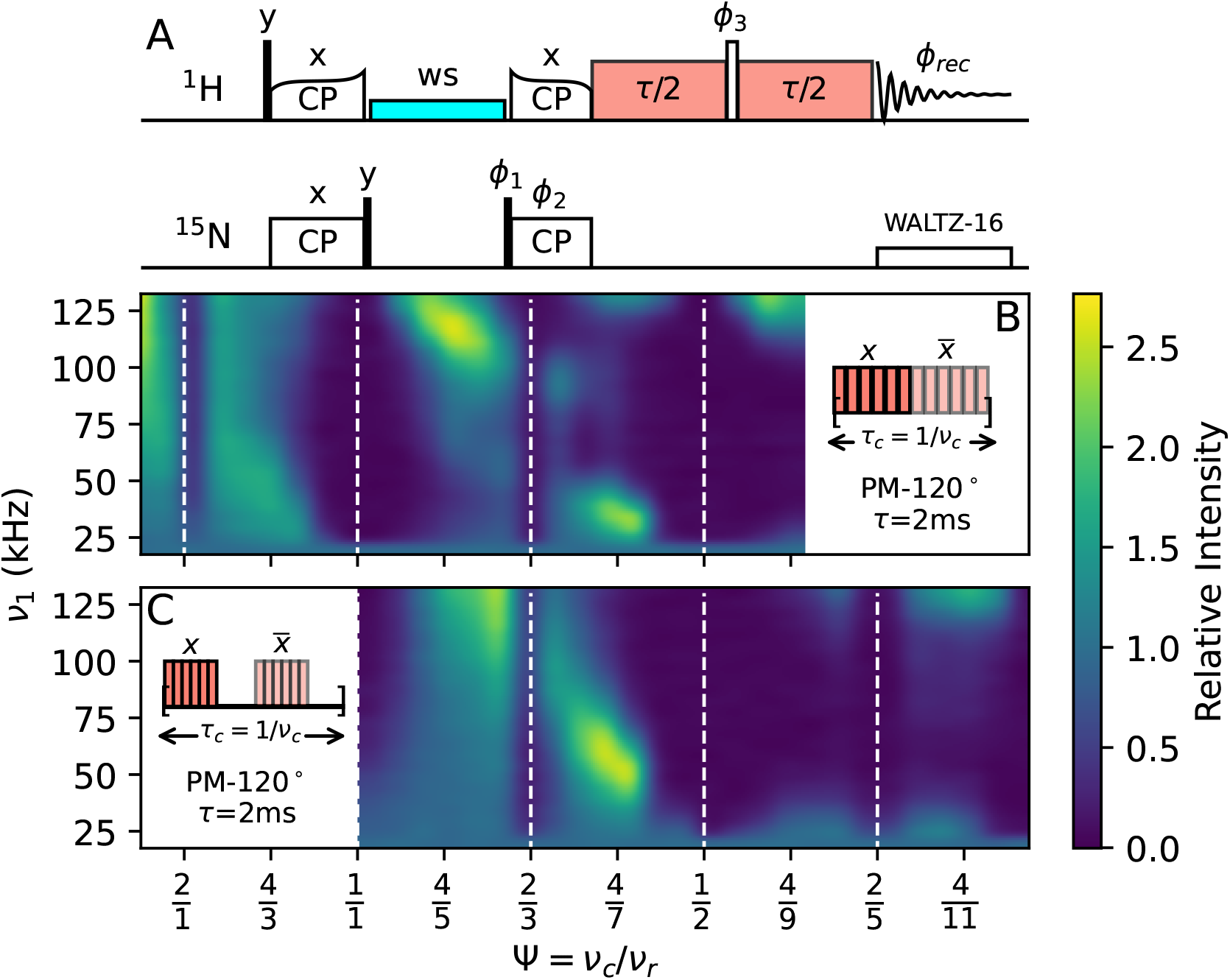
(A) Pulse sequence for the optimization of phase-modulated homonuclear decoupling using the spin-echo sequence. Thin, closed rectangles are *π/*2 pulses and open rectangles are *π*-pulses. The phases (divided by 90^*°*^) are as follows: *ϕ*_1_ = 31, *ϕ*_2_ = 0022, *ϕ*_3_ = 0000111122223333, *ϕ*_*rec*_ = 02202002. All experimental parameters are provided in the supporting information. (B, C) Representative experimental data for the optimization of homonuclear decoupling at the MAS frequency of 62.5 kHz on f-MLF, for the homonuclear decoupling sequence PM-120^*°*^(see Equation 2 for nomenclature). Ψ = *v*_*c*_*/v*_*r*_ and the rf amplitude are independently varied. The color-scale represents the intensity of the resonance observed with homonuclear decoupling after a 2.0 ms echo, normalized by the intensity in the absence of homonuclear decoupling for the same echo duration, for the leucine residue in f-MLF. A windowless homonuclear decoupling sequence is used for the plot in (B). Two windows (5.3 *μ*s each) are included per supercycle for the plot shown in (C). Theoretically predicted resonance conditions are indicated by dashed vertical lines. Optimization plots at different spinning frequencies and phase ramps are provided in Section S2 (Figures S1 to S4).

The starting magnetization in these experiments is prepared after two cross-polarization steps (Figure 1A), which are almost always required in multidimensional experiments on biomolecules. This leads to a partial selection of the sample that experiences a narrower distribution of *v*_1_ fields [44–47]. Figure S5 shows nutation curves for a sample of f-MLF in a single pulse and a ^15^N-filtered experiment based on cross-polarization. A significant improvement in rf-homogeneity is evident in these experiments. One can thus expect a better performance of the homonuclear decoupling sequence as compared to a single pulse, similar to the observations that demonstrate an improved performance of symmetry-based sequences in TOBSY recoupling after cross-polarization [48]. Alternate ways of achieving this such as restricted sample packing or the use of rf-selective pulses [28] were not considered here since they lead to a significant loss in sensitivity.

It is readily apparent from Figure 1B,C that only a few good homonuclear decoupling conditions exist in this range of rf amplitudes and decoupling cycle times, for both windowed and windowless sequences. For both 60 kHz, and 95 kHz spinning, two regions where homonuclear decoupling works well were identified: Ψ values between 0.52–0.63, and 0.70–0.78 (Figure 1, Figures S1 to S4). Both of these are close to the relatively narrow resonance condition corresponding to n=2, k=3 in Equation 1. Broad resonance conditions (n=k=1 and n=1, k=2) restrict the choice of Ψ to the above two regions, which in turn restricts the rf amplitude that can be used. The optimal decoupling conditions are thus fixed for each MAS frequency and can only coincidently occur at low rf amplitudes. The only way to allow the tailoring of these conditions to lower (or higher) rf amplitudes is to change the phase ramp of the homonuclear decoupling sequence while keeping its cycle time constant. This allows for the optimization of the rf amplitude independent of the cycle time. In principle, only the rf amplitude needs to be varied for each phase ramp with one of the two Ψ values identified above. However, for completeness, we report optimization plots that show the variation of all three parameters, i.e. the phase ramp, rf amplitude, and Ψ in Section S2 (Figures S1 to S4). Figure 2 shows that shallower phase ramps lead to a decrease in the optimal rf amplitude required for decoupling while these best decoupling conditions shift to higher rf amplitudes for a steeper phase ramp. Irrespective of the phase ramp used, the optimal region for homonuclear decoupling remain the same as above. The numerical value of the phase ramp itself has no meaning in the absence of any information about cycle time, but we have chosen the following nomenclature for a decoupling sequence consisting of 6 pulses for convenience in reporting these sequences:

**FIG. 2.**
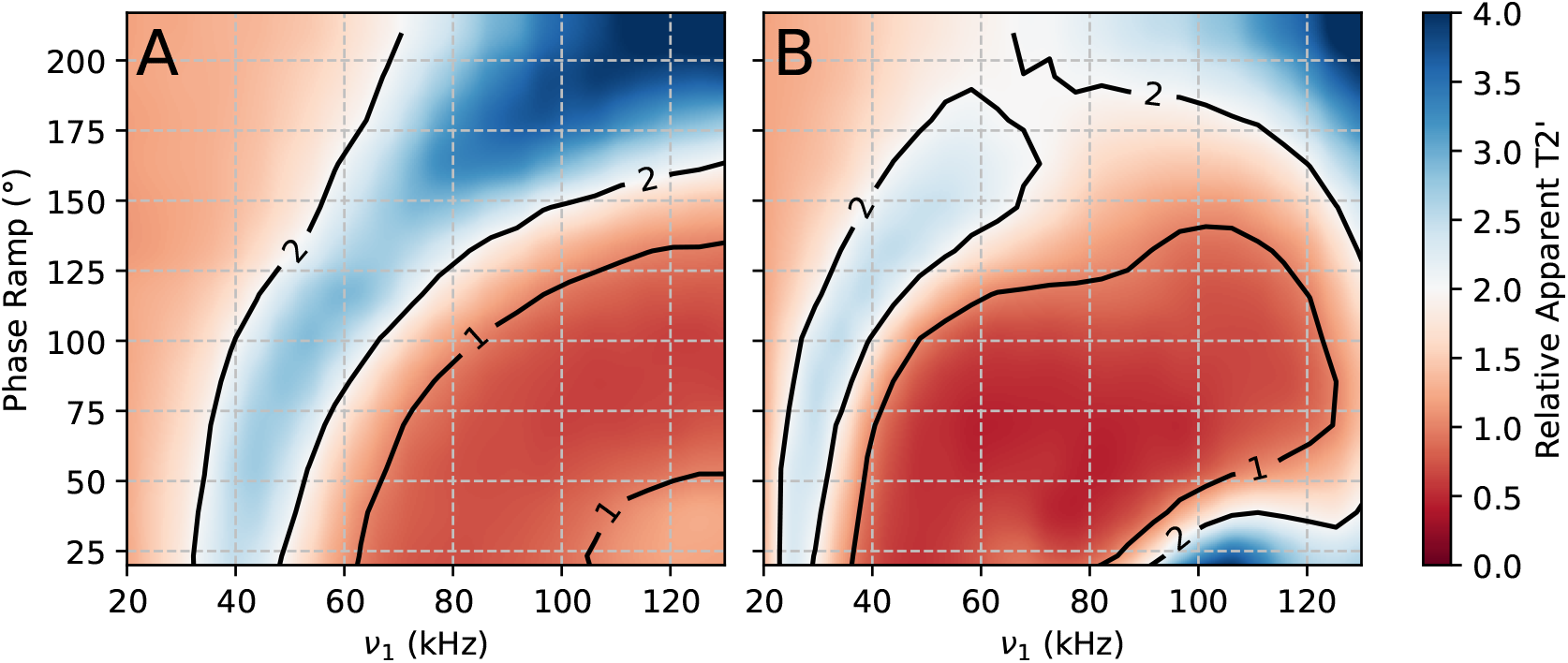
Improvement in apparent *T*_2_’ times (for L3 in f-MLF at 62.5 kHz MAS) at Ψ ∼ 0.57 for different phase ramps in a PM-*θ* sequence as a function of the rf amplitude used for homonuclear decoupling. (A) Using windowed homonuclear decoupling with a window duration of 5.3 *μ*s. (B) Using windowless homonuclear decoupling. Effective *T*_2_’ times which take into account the scaling factor, are reported in the Section S3.

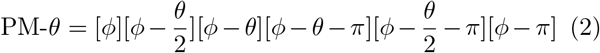

Here, *θ*, the phase ramp, is defined as the difference between the starting and ending phase of the sequence of 3 pulses, and *ϕ* is the starting phase. *ϕ* can be set to 0^*°*^, but we have chosen the value of ∼ 325^*°*^ to be consistent with the value that is used in the ‘m3m’ homonuclear decoupling shape that is available on Bruker spectrometers. Note that this sequence consists of two linear phase ramps of 3 pulses each, and corresponds to the frequency-switched version of the Lee-Goldburg experiment. The two-step supercycle is applied on top of this sequence as given by Equation 3:

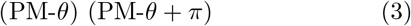

Experimental datasets in Figure 1B,C correspond to the supercycled version of PM-120^*°*^. With this setup, we now discuss the improvements in coherence lifetimes and resolution that are obtained when each sequence PM*θ* is individually optimized, and determine the optimal setup for achieving the highest resolution at the MAS frequencies of *∼*60 kHz and *∼*95 kHz.

### A. Improvement in transverse coherence decay times for ^1^H

^1^H transverse relaxation times were measured in a spin-echo experiment using a mono-exponential fit (*T*_2_’, Section S3). These are proportional to the MAS frequency in absence of homonuclear decoupling, as homogeneous contributions dominate relaxation in protons under these conditions in non-deuterated samples [5, 10, 11]. While the assumption of a mono-exponential decay is not strictly correct for powder samples under MAS, it is a convenient description to estimate the efficiency of the sequences described here. For amide protons, *T*_2_^*′*^ increases from *<*1.0 ms below the spinning frequency of 40 kHz to 3.0–5.0 ms at the spinning frequency of 100 kHz and further to 5.0–8.5 ms at 160 kHz spinning [12, 13]. These numbers are consistent with those observed for the samples used in our study (Section S3). The relatively fast decay of magnetization precludes the use of pulse sequences that rely on long refocusing blocks on ^1^H under MAS (for example, INEPT [49]). The optimization procedure described in the previous section results in PM-*θ* parameters that achieve the longest *T*_2_’ times. A substantial increase in the *T*_2_’ times by a factor of *∼* 2–4 is seen at the spinning frequencies of 62.50 kHz and 95.24 kHz compared to MAS alone (Figure 2, Section S3). As expected, the optimal rf increases with increasing phase ramp (Figure 2). For the windowed sequences, a high scaling factor (*>*0.8) is observed for most of the conditions (Tables S2 and S5). This is not always the case for windowless sequences, where significantly lower scaling factors were observed (Tables S3 and S6). The refocused-INEPT (rINEPT) experiment is a stringent test for demonstrating these improvements as its efficiency depends primarily on ^1^H 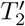. An ^15^N-filtered ^1^H 1D spectrum for f-MLF was acquired using a windowless homonuclear decoupling sequence during the INEPT-transfer period on ^1^H (Figure S13). Efficiencies of ∼ 25% were seen for residues L2 and F3 in f-MLF, and ∼ 60% for M1 due to the significantly higher *T*_2_’ for this residue. These numbers are consistent with estimates based on the one-bond scalar coupling of 92 Hz for an amide bond for the observed effective *T*_2_’ times. As expected, the selectivity of rINEPT-based transfer from ^15^N to ^1^H is higher than cross-polarization (CP), which showed transfers to aliphatic protons even with reduced contact times (Figure S13). Applications that require highly selective transfers can take advantage of the higher efficiency of CP for the initial transfer from ^1^H to ^15^N and use rINEPT for the reverse transfer. Figure S13 shows that this boosts the relative sensitivity to ∼ 55% for L2 and F3 (84% for M1) in f-MLF. A similar relative efficiency of 25% was observed for the rINEPT experiment in a sample of GB1. 2D spectra acquired using CP and rINEPT show a nearly identical set of peaks (Figure S14). While the general application of rINEPT-based experiments is not recommended due to its lower sensitivity even with the approach described here, we anticipate a direct application of these sequences to other experiments that require a long transverse relaxation time during a spin-echo sequence on ^1^H [50–52]. This increase in *T*_2_’ time also bodes well for obtaining a substantial improvement in the detected linewidths in the direct dimensions, as long as the deleterious effects of rf-inhomogeneity are avoided. The following sections discuss the condition under which improved resolution was obtained.

### B. Improved resolution with windowed detection

Most of the multidimensional experiments used in biomolecular solid-state NMR consist of a single ^1^H dimension (the direct dimension). This is due to a combination of its high sensitivity and poor chemical-shift dispersion as compared to ^13^C and ^15^N, which are instead evolved in the indirect dimensions. Hence, we emphasize the detection of ^1^H in the direct dimension in this article, while noting that these experiments can also be done when ^1^H is evolved in the indirect dimension. This approach requires that homonuclear decoupling sequences are used together with windowed acquisition. The optimization procedure described above can be adapted directly to achieve this by including the window as a part of the homonuclear decoupling sequence (Figure 1B). The duration of the window for detection depends on the probe being used, and was optimized to 5.3-5.4 *μ*s for the 1.3 mm and 0.7 mm probes used here. Note that the window itself occupies a substantial portion of the rotor period at these spinning frequencies (∼ 33% at 62.50 kHz and ∼ 50% at 95.2 kHz without homonuclear decoupling). We see an increase in the rf requirement with the incorporation of a window (Figure 1B,C, Figure 2), but as expected, the optimal Ψ values for decoupling remain similar to the case without the window. Of these, we chose to work with conditions that have a low rf amplitude requirement (Figure 1, Figures S1 to S4). This not only avoids increasing MAS-modulated rf-inhomogeneity, but is also the prudent choice keeping in mind the need to minimize rf-induced heating in these samples. Under these conditions, a striking improvement in resolution is seen at both 62.50 kHz and 95.24 kHz for a sample of f-MLF, as shown in Figure 3A,B. Here, 13 kHz rCW^ApA^ heteronuclear decoupling was used on ^15^N [53, 54]. For f-MLF, optimization of heteronuclear decoupling is straightforward as the resolution for the three observed resonances can be directly measured under different heteronuclear decoupling sequences [32]. rCW^ApA^ decoupling was used as it can accommodate large variations in the cycle frequency at a constant rf amplitude by changing the length of continuous-wave (CW) block. The spectra without homonuclear decoupling were obtained in the presence of standard WALTZ16 decoupling at rf amplitude of 13 kHz. The full width at half maximum (fwhm) reduces by a factor of ∼ 1.6 for L2 and F3 and a factor of ∼ 1.3 for the narrower M1 resonance Figure 3A. This improvement continues at the spinning frequency of 95.24 kHz, where improvements by a factor of 1.6, 1.2, and 1.4 are seen for the three resonances (Figure 3B). With these experiments on f-MLF, we draw three conclusions: (i) The relative improvement is dictated by the overall broadening that the resonances experience, i.e., broader resonances benefit to a higher degree from the application of homonuclear decoupling (ii) Spectra obtained using homonuclear decoupling at 62.50 kHz MAS can approach or even improve upon the resolution obtained at 95.24 kHz without homonuclear decoupling (Figure 3A,B), and (iii) Spectral resolution continues to improve at 95.24 kHz upon the application of simultaneous homoand heteronuclear decoupling (Figure 3B). With these results in hand, we turn our attention to larger proteins systems, which are a stringent test for the improvement obtainable with these methods.

**FIG. 3.**
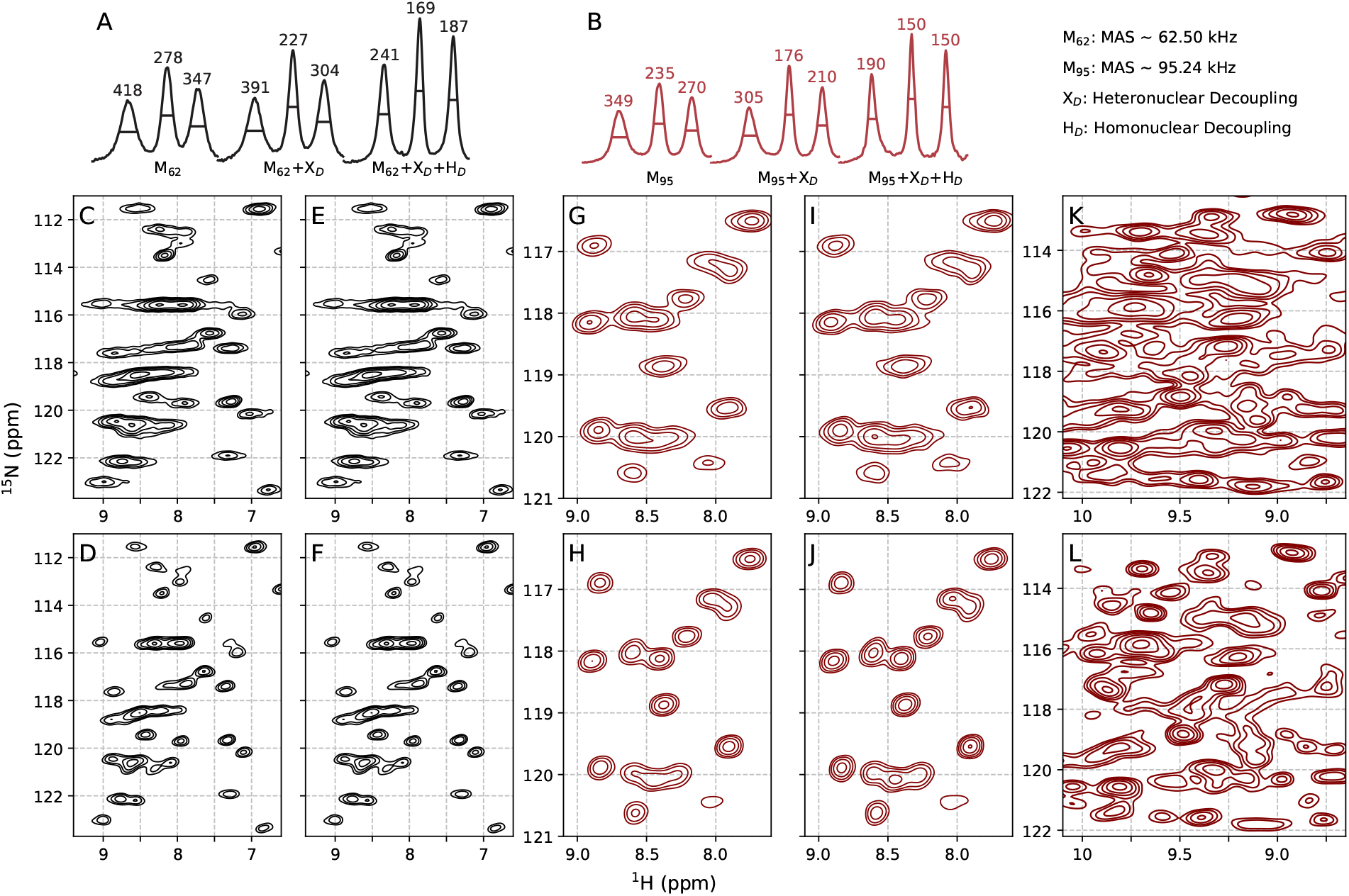
^15^N-^1^H spectra of f-MLF (A, B), microcrystalline GB1 (C-J), and ParM filaments (K, L) at the MAS frequencies of 60 kHz (black) and 95 kHz (red) that demonstrate the Line narrowing obtained using homonuclear decoupling and windowed detection in the direct dimension. ^15^N-filtered ^1^H 1D spectra for f-MLF without homo- and hetero-nuclear decoupling (left), with heteronuclear decoupling (middle), and with both, homo- and heteronuclear decoupling (right) at *v*_*r*_ =62.50 kHz (A) and 95.24 kHz (B) show that the linewidths at half-maximum for each peak improve by a factor of 1.2-1.6, with the larger improvements seen for the broader peaks. All spectra in A, B were acquired with windowed detection for a direct comparison. (C-F) Comparison between the spectra of GB1 obtained at *v*_*r*_ =60.61 kHz without (C, E) and with homonuclear decoupling (D, F). Spectra in E, F have the same underlying dataset as in C, D except that they are processed with an additional window function in the ^1^H dimension. (G-I) Comparison between the spectra of GB1 obtained at *v*_*r*_ =95.24 kHz without (G, I) and with homonuclear decoupling (H, J). Spectra in I, J have the same underlying dataset as in G, H except that they are processed with an additional window function in the ^1^H dimension. (K, L) Comparison between the spectra of ParM obtained at *v*_*r*_ =95.24 kHz without (K) and with homonuclear decoupling (L). All 2D spectra are plotted with contour levels spaced by a multiplicative factor of 1.4 starting with a contour level at 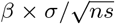, where *σ* is the noise rmsd, *ns* is the number of scans, and *β* is 20 (C-F), 12 (G-J), and 6 (K-L). All spectra for f-MLF were acquired with windowed detection. Spectra without homonuclear decoupling were acquired with standard detection for GB1 and ParM. Experimental parameters and complete spectra for all three samples under different conditions are reported in Section S4.

In microcrystalline GB1 (non-deuterated) at 60.61 kHz, a substantial improvement in resolution is seen, as shown in Figure 3C-F. The average linewidth in this system decreases from 210 ±39 Hz to 148± 32 Hz, and the ratio of linewidths with and without homonuclear decoupling is 0.7 ±0.1 (Figure 4A-C). The improvement in resolution is correlated with the linewidth of the peak in question. Broader peaks show a significantly higher improvement than the peaks that are already narrow in absence of homonuclear decoupling (Figure 4C). Some of the narrowest peaks show a higher fwhm due to a combination of a relatively small improvement due to homonuclear decoupling and a loss in effective resolution due to the chemical-shift scaling factor. Nevertheless, most of the peaks show a substantial improvement in linewidths, with some of the broadest peaks showing linewidths that are narrower by a factor of ∼2. Similar results are obtained in a sample of GB1 at 95.24 kHz spinning. The average fwhm decreases from 147± 33 Hz to 103 18 Hz, with a similar ratio of linewidths with and without homonuclear decoupling (0.7± 0.2, Figure 3G-J and Figure 4D-F). Similar to f-MLF, we see that the average linewidth at 60 kHz spinning (148*±*31 Hz) in the presence of homonuclear decoupling is close to the linewidths observed without homonuclear decoupling at 95 kHz spinning (147 ± 32 Hz). Even at the higher spinning frequency, the highest improvement in broad peaks, while narrower peaks tend to to show reduced improvement (Figure 4F).

**FIG. 4.**
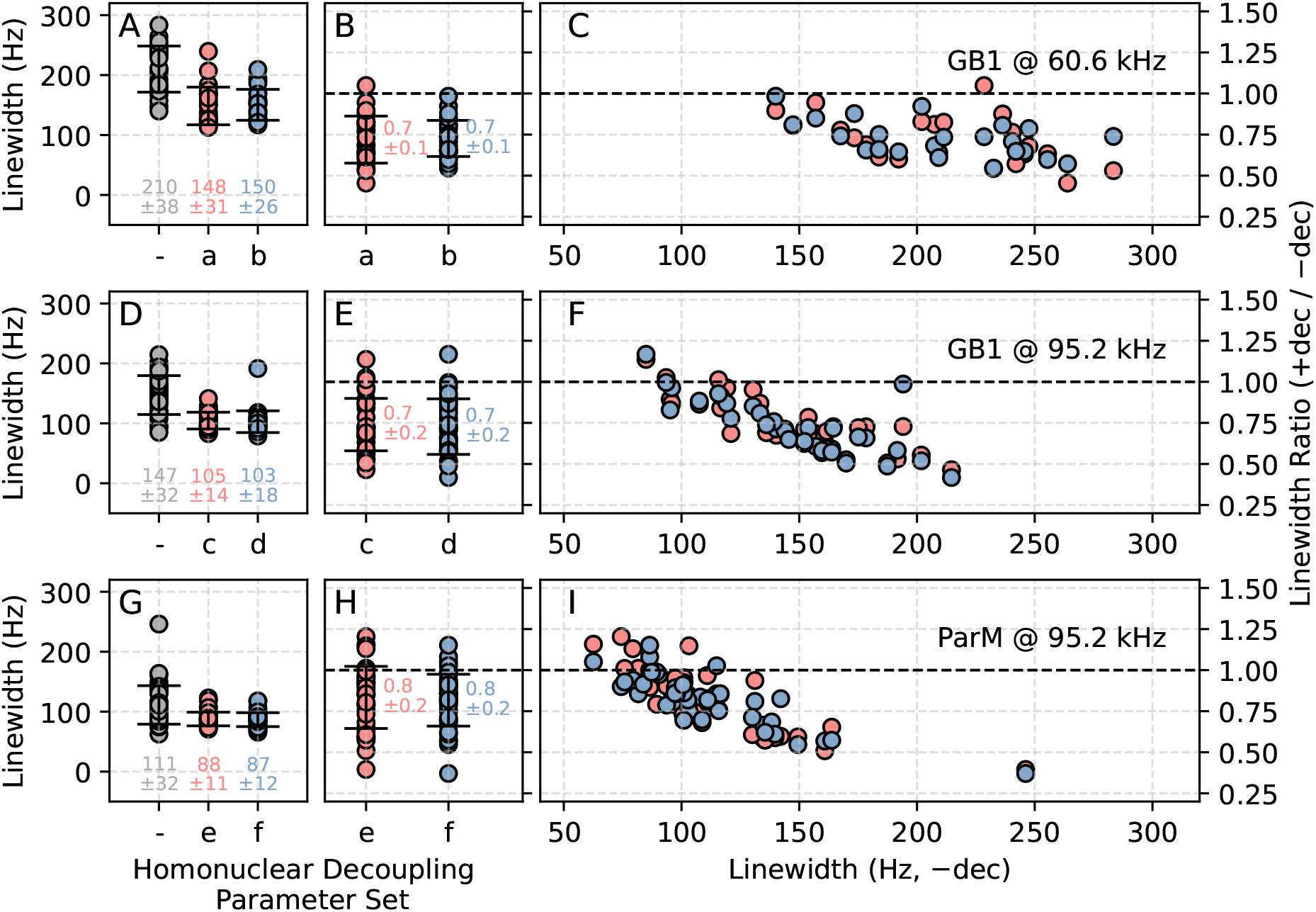
Analysis of site specific improvements in line widths due to homonuclear decoupling for GB1 (A-F) and ParM (G-I) at the MAS spinning frequencies of 60.61 kHz (A-C) and 95.24 kHz (D-I). Peaks that were resolved in the 2D spectra were selected for this analysis. (A, D, G) show that the a significant reduction the the average linewidth upon homonuclear decoupling at two conditions. (B, E, H) show that the ratio of the linewidths with and without homonuclear decoupling spans a range from 0.5-1.2, with a majority of the peaks showing lower linewidths in the presence of homonuclear decoupling. (C, F, I) show that the improvement in linewidth is correlated to the measure linewidth of the resonances without homonuclear decoupling. The largest reduction in linewidth is seen for peaks that are very broad, while peaks that are already narrow in the absence of homonuclear decoupling tend to show a reduced improvement in the linewidth. For some of the narrowest peaks, there is an effective broadening due to homonuclear decoupling that can be attributed to smaller improvements and a net broadening due to the scaling factor (0.80-0.88). The homonuclear decoupling parameter sets used were as follows: **a**: PM-101^*°*^Ψ=0.58 rf=45 kHz; **b**: PM-62^*°*^Ψ=0.58 rf=40 kHz; **c**: PM-120^*°*^Ψ=0.55 rf=124 kHz; **d**: PM-101^*°*^Ψ=0.55 rf=124 kHz; **e**: PM-62^*°*^Ψ=0.55 rf=82 kHz; **f** =PM-120^*°*^Ψ=0.55 rf=124 kHz.

For larger proteins, a high density of resonances makes identification of resonances challenging even when they are individually narrow. The number of resolved peaks reduces rapidly in the presence of few broad resonances which can overlap with several peaks. This is clearly seen in a 2D ^15^N-^1^H HSQC spectrum of a prokaryotic cytoskeletal protein ParM, a protein consisting of 326 amino acids prepared in a filamentous state (F-state) by the addition of a non-hydrolyzable ATP analogue ATP*γ*S (Figure 3K). ParM in its F-state forms highly homogeneous filaments that result in spectra with the average linewidth approaching 110 Hz in absence of homonuclear decoupling at the spinning speed of 95.24 kHz (based on 41 peaks that are resolved in the 2D ^15^N-^1^H spectrum). As such it represents a very challenging system to further improve upon, as the inherent linewidths are narrower than even microcrystalline GB1. As shown in Figure 3L, a significant improvement in the resolution is seen in the ^15^N-^1^H correlation spectrum when acquired with windowed detection and homonuclear decoupling. We are able to estimate the improvement in the resolution by analyzing 41 peaks which are resolved in both the spectra. The average line width decreases from 111 ± 33 Hz to 87 ± 12 Hz, showing an average improvement of 0.8 ± 0.2 Figure 4G-H. Similar to the case of f-MLF and GB1, it is the broadest peaks that show the highest improvement, which explains the dramatically improved resolution seen in the central region of the spectrum with a high density of the resonances (Figure 4I).

Note that, unlike f-MLF, where heteronuclear decoupling was optimized by monitoring the resonance fwhm directly, heteronuclear decoupling was optimized for GB1 and ParM by measuring the effective *T*_2_^*′*^ as a function of the cycle frequency of the rCW^ApA^ sequence, greatly simplifying the above experiments (Section S5). This improved resolution will be of critical importance in studying large complexes such as ParM, where experiments are challenging in spite of the high resolution in the ^13^C and ^15^N. We expect this improved resolution to not only simplify assignments but also enable the use of 2D or 3D experiments to obtain site-specific information on dynamics in these complexes.

### C. Signal-to-noise ratios in windowed detection

Only a part of the signal is sampled during windowed detection. Not accounting for any improvements due to narrower resonances, one can expect a maximum sensitivity that is proportional to the square root of the fraction of the signal that is acquired in these experiments ([26, 55]). Additional factors such as an increased noise floor can further reduce the sensitivity ([26]). At the conditions described above, we estimate a theoretical maximum sensitivity of ∼0.47 and ∼0.57 for 1.3 mm and 0.7 mm probes, respectively. The actual sensitivity, as determined by comparing the signal-to-noise ratio in a standard detection (Figure 5A,D) to that acquired in a windowed detection experiment without homonuclear decoupling (Figure 5B,E), was found to be∼ 0.42 and ∼0.31 for two probes respectively. These results are in line with those obtained at lower MAS frequencies of 5-28 kHz where an increase in the noise floor of∼ 1.5-1.9 was observed [26]. We note that different probes have a different increase in the noise floor; the 1.3 mm HCN probe has a relatively small increase in the noise floor (1.12) as compared to the 0.7 mm probe, where this increase was estimated to be 1.83. The experiments are hence, significantly more sensitive on the 1.3 mm probe. It is at present unclear what parameters in the construction of the probes influences these, and we have not attempted to address this loss here.

**FIG. 5.**
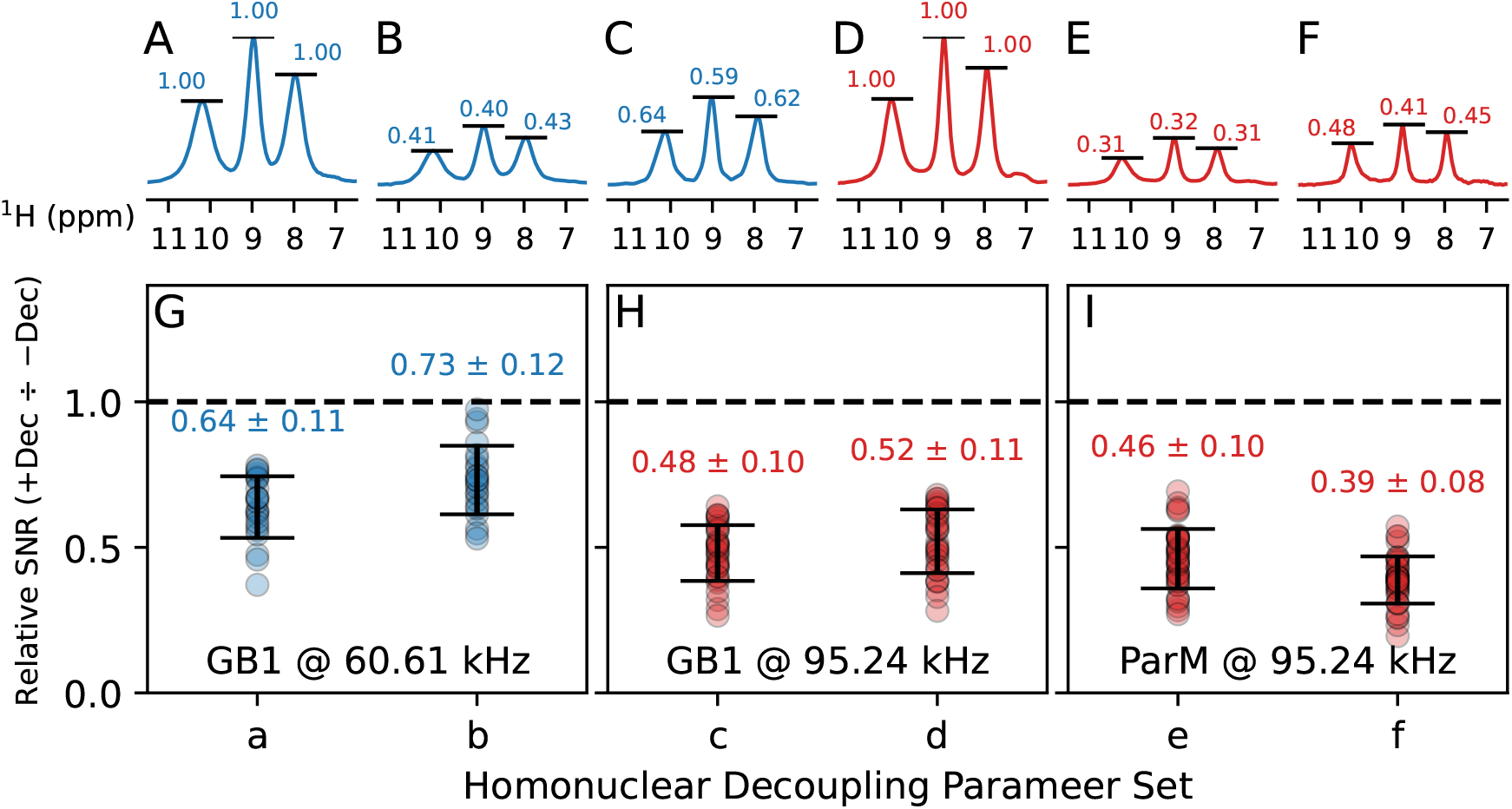
Comparison of signal-to-noise ratios (SNR) obtained using window and standard detection for f-MLF (A-F), GB1 (G-H) and ParM (I) at the MAS frequencies of 60-62.5 kHz (blue) and 95.24 kHz (red). The efficiency of windowed detection, as seen from the data on f-MLF on our probes at the MAS frequency of 60.61 kHz is 0.41 (B), while that at 95.24 kHz is 0.31 (E). Line narrowing due to homonuclear decoupling results in improvement in the SNR by a factor proportional to the reduction the in the linewidth (1.5-1.6), resulting in a relative sensitivity of 0.62 and 0.45 with respect to standard detection. The average relative SNR for the resolved peaks in GB1 at 60.61 kHz spinning is 0.73 wheres that for GB1 and ParM at 95.24 kHz spinning is 0.52 and 0.46, respectively. Note that the SNR here is defined as the peak height divided by the noise rmsd, and not the peak integral. Homonuclear decoupling parameter sets **a-f** are described in the caption for Figure 4. 1D datasets for f-MLF, GB1, and ParM at different homonuclear decoupling conditions are reported in Section S6.

The application of homonuclear decoupling improves resolution in some cases by a factor of 1.5 to 2.0 (Figure 4). One can expect higher sensitivity for these resonances as a direct result of this line narrowing (sensitivity is defined as the peak height divided by noise rmsd). The improved resolution in f-MLF boosts the sensitivity by almost 50%, from approximately 0.42 to 0.60 at the spinning frequency of 62.50 kHz (Figure 5C).

A similar 50% improvement in sensitivity is also seen at the higher spinning frequency of 95.24 kHz where it goes from 0.31 to 0.45 (Figure 5F). As expected, this improvement is highest for the resonance that is broad (L2) due to the greater reduction in the linewidth. These numbers translate well to the proteins GB1 and ParM at the two spinning frequencies. GB1 at the spinning frequency of 60.61 kHz shows an average sensitivity of 0.73 ± 0.12 as compared to standard detection Figure 5G. A combination of substantially narrower resonances and better probe performance compensates for the loss in sensitivity under these spinning conditions. As expected from the data on f-MLF, the sensitivity for the GB1 and ParM samples at 95.24 kHz is 0.52 ± 0.11 and 0.46 ± 0.10, respectively (Figure 5G-I).

We note here that the application of homonuclear decoupling during ^1^H evolution in the indirect dimension does not result in reduced sensitivity. Experiments such as 4D-HN(COCA)NH [56], where ^1^H is evolved in both, the direct and indirect dimensions, will benefit to a greater degree from the application of these sequences. For detection in the direct dimension, the loss in sensitivity is comparable to that associated with performing experiments on probes with lower rotor volumes. For example, a 0.7 mm probe is *∼*0.4*×* the sensitivity of a 1.3 mm probe [11], while a 0.4 mm probe is *∼*0.33*×* the sensitivity of a 0.7 mm probe [13]. The improved resolution obtained using homonuclear decoupling on probes that accommodate higher volume rotors can thus allow experiments to be done with both higher resolution and sensitivity as compared to similar experiments on a lower volume probe. For our 1.3 mm probe, a relatively small increase in the noise floor together with a large improvement in resolution nearly compensates for the loss in sensitivity, making the utility of these experiments apparent in nearly all cases. At the higher MAS frequencies afforded by the 0.7 mm probe, we expect that the substantially improved resolution will be critical when tackling large complexes with a very high density of resonances. Although we expect that the *T*_2_’ times in the presence of homonuclear decoupling will continue to increase at even higher MAS frequencies, the resolution at these MAS frequencies will ultimately be limited by inhomogeneous broadening, which is expected to become the dominant factor at MAS frequencies approaching 200 kHz [11, 13]. We also note that higher MAS frequencies result in longer transverse relaxation times for ^15^N and ^13^C nuclei, and are expected to lead to higher transfer efficiencies in multidimensional experiments. Nevertheless, homonuclear decoupling together with windowed acquisition will be essential to maximise the information content in spectra recorded at spinning frequencies between 60-100 kHz, which are now commonly available.

## IV. CONCLUSIONS

We have shown here that spectra with significantly improved resolution over MAS alone can be obtained by the application of homonuclear decoupling sequences coupled with windowed detection at fast MAS frequencies. These sequences are easy to optimize and have a relatively low rf requirement. A substantial improvement is demonstrated on standard samples of f-MLF and the microcrystalline protein GB1 as well as a filamentous preparation of a cytoskeletal protein ParM. These sequences have a relative sensitivity of 0.4-0.7 as compared to standard detection. The improved resolution that these sequences enable will be of importance when working with large complexes where the high density of broad resonances makes site-specific analysis challenging. We anticipate a direct application of these sequences at MAS frequencies between 60-100 kHz, where one can take advantage of the comparatively higher rotor volumes and still achieve resolution that would have otherwise required experiments with rotors of a substantially smaller volume, and hence, lower sensitivity.

## V. METHODS

Detailed description of the preparation of samples, parameters used in NMR experiments, and pulse sequence codes for Bruker spectrometers is provided in the supporting information. Pulse sequences associated with this article are also available for download at https://github.com/kaustubhmote/pulseseq.

## VI. ACKNOWLEDGEMENTS

The authors acknowledge intramural funds at TIFR Hyderabad from the Department of Atomic Energy (DAE), India, under Project RTI 4007. We thank Profs. Matthias Ernst and Vipin Agarwal for discussions.

