## Supporting Information for "Beyond Simply Spinning: Improving ^1^H Resolution at Fast Magic-Angle-Spinning Frequencies Using Combined Rotation and Multiple Pulse Spectroscopy"

### S1. METHODS AND MATERIALS

#### A. U-$^{1}$H,$^{13}$C,$^{15}$N f-MLF

Uniformly $^{13}$C and $^{15}$N labeled f-MLF was purchased from Cambridge Isotopes Limited, as used as is. Samples were fully packed in 1.3 mm and 0.7 mm zirconia rotors (Bruker) without any spacers. All optimizations were performed on uniformly $^{13}$C and $^{15}$N labeled tripeptide N-formyl-methionyl-leucyl-phenylalanine (f-MLF) at MAS frequencies of 62.50 and 95.24 kHz on an Avance-III Bruker 700 MHz (16.5 T) spectrometer equipped with three-channel 1.3 mm (HCN) and 0.7 mm (HCN) probes and Topspin 3.5pl7. A spin echo experiment with a constant echo duration of 2 ms and 3 ms, respectively, was employed to compare the efficiency of different homonuclear decoupling schemes in terms of $T_2'$ at 62.50 and 95.24 kHz, respectively. Different phase ramps were used ranging from 20 to 240 degrees as described in the main text. RF amplitude was varied from 20 to 130 kHz and 20 to 225 kHz at 62.50 and 95.24 kHz, respectively. Phase-ramped decoupling was implemented with each phase-ramp consisting of 3 phases varied linearly The total supercycled decoupling block thus consists of 12 pulses with a constant amplitude and varying phases. The length of each pulse was varied between 0.5-3.5 $\mu$s in 0.1 $\mu$s steps. The duration of the window was optimized in the presence of homonuclear decoupling to get the maximum signal-to-noise ratio and to minimize any artifacts. A value of 5.3 $\mu$s was found to be optimal for both the probes. The cycle time reported is for the supercycled block. For the windowed decoupling schemes, the reported cycle time includes the duration of the window. All spectra with homonuclear decoupling and windowed acquisition are scaled by the experimentally determined scaling factors. Scaling factors for amide protons in f-MLF for non-windowed decoupling sequences were determined by recording $^{1}$H-$^{1}$H correlation after $^{15}$N filtering, with homonuclear decoupling being applied during indirect $^{1}$H evolution.

### B. U-$^{1}$H,$^{13}$C,$^{15}$N GB1

Microcrystalline GB1 was prepared using previously described protocols [1, 2]. Experiments on GB1 were performed at the MAS frequencies of 60.61 kHz and 95.24 kHz using the same spectrometer as for f-MLF, with the same MAS probes. The temperature Of the inlet gas was set to -25°C, which resulted in the actual temperature of approximately +10°C, as estimated by the chemical shift of the water resonance.

#### C. U-$^{1}$H,$^{13}$C,$^{15}$N ParM

Uniformly $^{13}$C and $^{15}$N labeled protein was as an N-terminal 6x-His tagged construct in a pDEST-based vector [3]. M9 medium suppemented with 2g/L $^{13}$C-glucose and 1g/L $^{15}$N-ammonium chloride was used to uniformly label the sample for NMR experiments. Protein expression was induced with 1.0 mM IPTG in Rosetta(DE3) cells. The polyhistidine tag was cleaved using TEV protease after Ni-affinity based purification, and the protein further purified to homogeneity using size-exclusion chromatography. Filaments in the F-state were prepared by the addition of a non-hydrolyzable ATP analog ATP-$\gamma$S to a solution of 4 mg/mL WT-ParM. The polymerization buffer contained 25 mM Tris, 20 mM KCl, 5 mM $MgCl_2$, 2 mM DTT, and 5% glycerol at pH 7.6. Experiments on these filaments were done at the MAS frequency of 95.24 kHz on the same spectrometer and probe described above. The temperature

*

of the sample was regulated at +18°C (as estimated by the chemical-shift of the water resonance) by setting the temperature of the inlet gas to -25°C to compensate for friction-induced heating.

### D. Experimental details

TABLE S1. Experimental parameters for 2D experiments that are common for all experiments.

| | MLF | MLF | GB1 | GB1 | ParM |
|---|---|---|---|---|---|
| MAS (kHz) | 62.50 | 95.24 | 60.61 | 95.24 | 95.24 |
| Temperature (Set) | 1°C | -25°C | -25°C | -25°C | -25°C |
| Temperature (Estimated at sample) | 40°C | 15°C | 15°C | 18°C | 18°C |
| CP $^1$H $\rightarrow$ $^{15}$N ($^1$H, kHz) | 125.8 | 88 | 93.5 | 86 | 86 |
| CP $^1$H $\rightarrow$ $^{15}$N ($^{15}$N, kHz) | 39.4 | 23 | 23 | 23.6 | 20 |
| CP $^{15}$N $\rightarrow$ $^1$H ($^1$H, kHz) | 116.2 | 83 | 90.5 | 79 | 86 |
| CP $^{15}$N $\rightarrow$ $^1$H ($^{15}$N, kHz) | 40.4 | 23 | 23 | 23.6 | 20 |
| CP $^1$H $\rightarrow$ $^{15}$N ($t$, $\mu$s) | 1600 | 1000 | 1000 | 1000 | 900 |
| CP $^{15}$N $\rightarrow$ $^1$H ($t$, $\mu$s) | 400 | 1000 | 800 | 1000 | 900 |
| Recycle Delay (s) | 1.0 | 1.0 | 1.0 | 1.0 | 0.89 |
| $^1$H decoupling (sequence) | rCW$^{\mathrm{ApA}}$ | rCW$^{\mathrm{ApA}}$ | rCW$^{\mathrm{ApA}}$ | rCW$^{\mathrm{ApA}}$ | rCW$^{\mathrm{ApA}}$ |
| $^1$H decoupling (kHz) | 11 | 11 | 11 | 11 | 11 |

### S2. OPTIMIZATION OF HOMONUCLEAR DECOUPLING

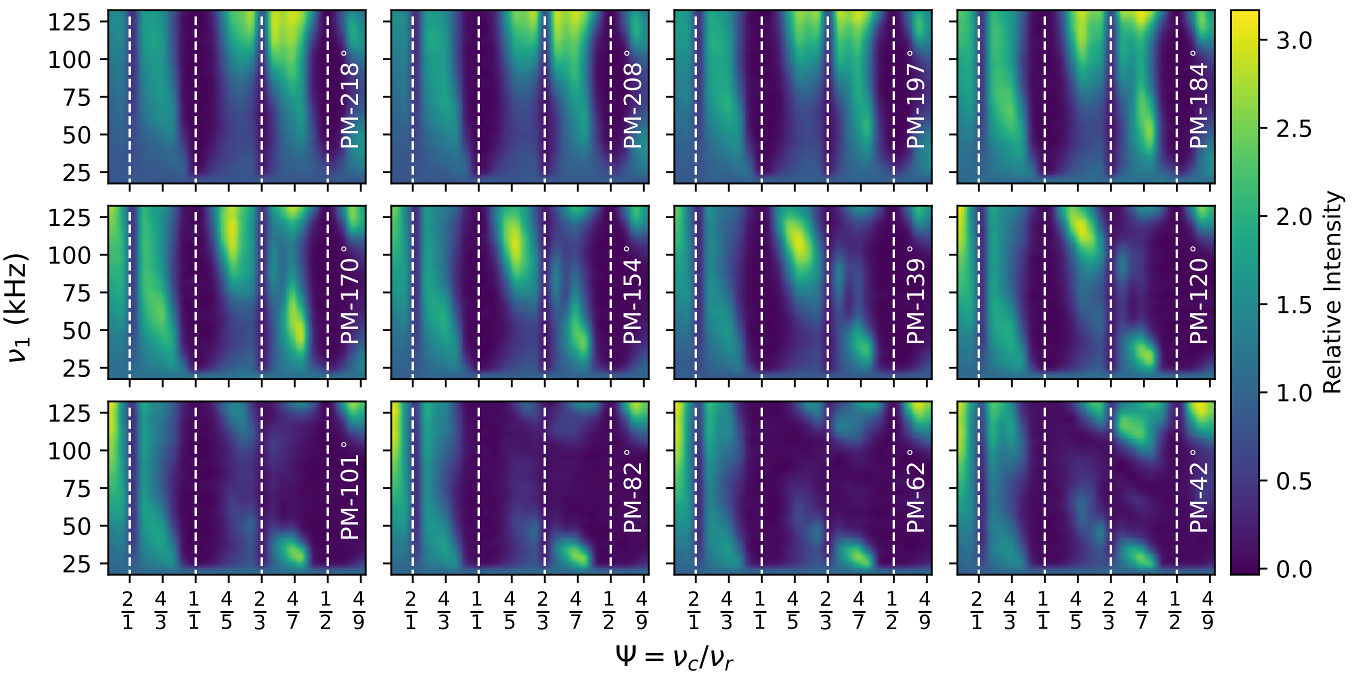


FIG. S1. Experimental data for the optimization of homonuclear decoupling at the MAS frequency of 62.50 kHz on a sample of the tripeptide f-MLF, for the homonuclear decoupling sequence PM-$\theta$, as indicated for each subplot (see main text for the nomenclature). $\Psi = \nu_c/\nu_r$ and the rf-amplitude are independently varied. The color-scale represents the intensity of the resonance observed with homonuclear decoupling after a 2.0 ms echo, normalized by the intensity in the absence of homonuclear decoupling for the same echo duration, for the leucine residue in f-MLF. A windowless version of PM-$\theta$ with a 3-pulse phase ramp is used. Theoretically predicted resonance conditions are indicated by dashed vertical lines.

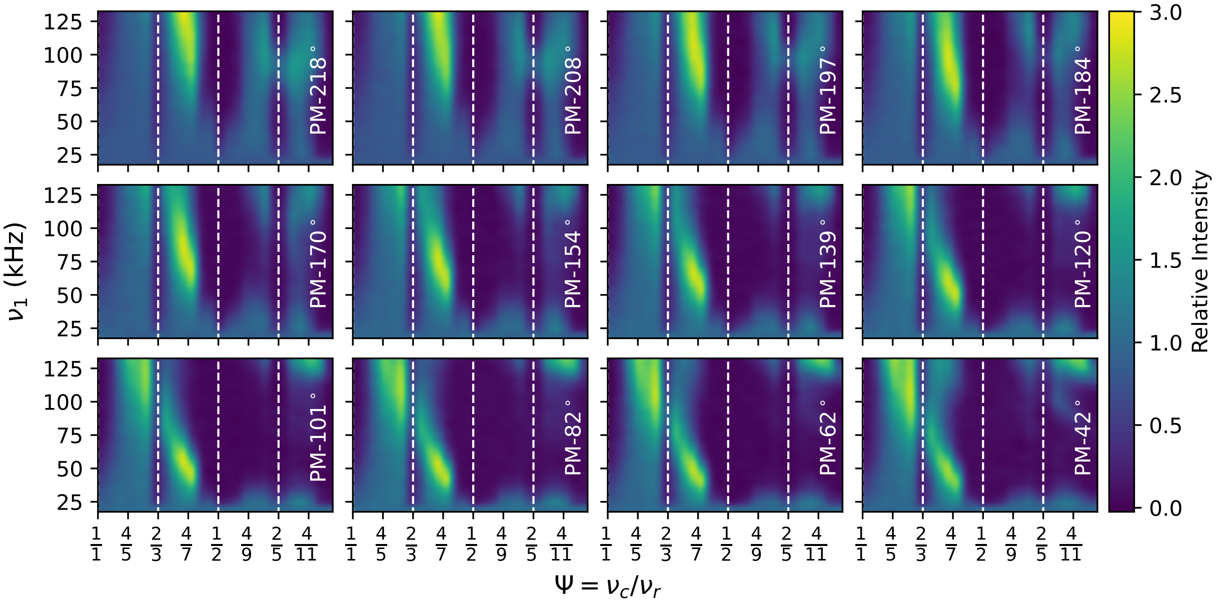


FIG. S2. Experimental data for the optimization of homonuclear decoupling at the MAS frequency of 62.50 kHz on a sample of the tripeptide f-MLF, for the homonuclear decoupling sequence PM-$\theta$, as indicated for each subplot (see main text for the nomenclature). $\Psi = \nu_c/\nu_r$ and the rf-amplitude are independently varied. The color-scale represents the intensity of the resonance observed with homonuclear decoupling after a 2.0 ms echo, normalized by the intensity in the absence of homonuclear decoupling for the same echo duration, for the leucine residue in f-MLF. A windowed version of PM-$\theta$ with two 5.3 $\mu$s windows per supercycle and a 3-pulse phase ramp is used for these plots. Theoretically predicted resonance conditions are indicated by dashed vertical lines.

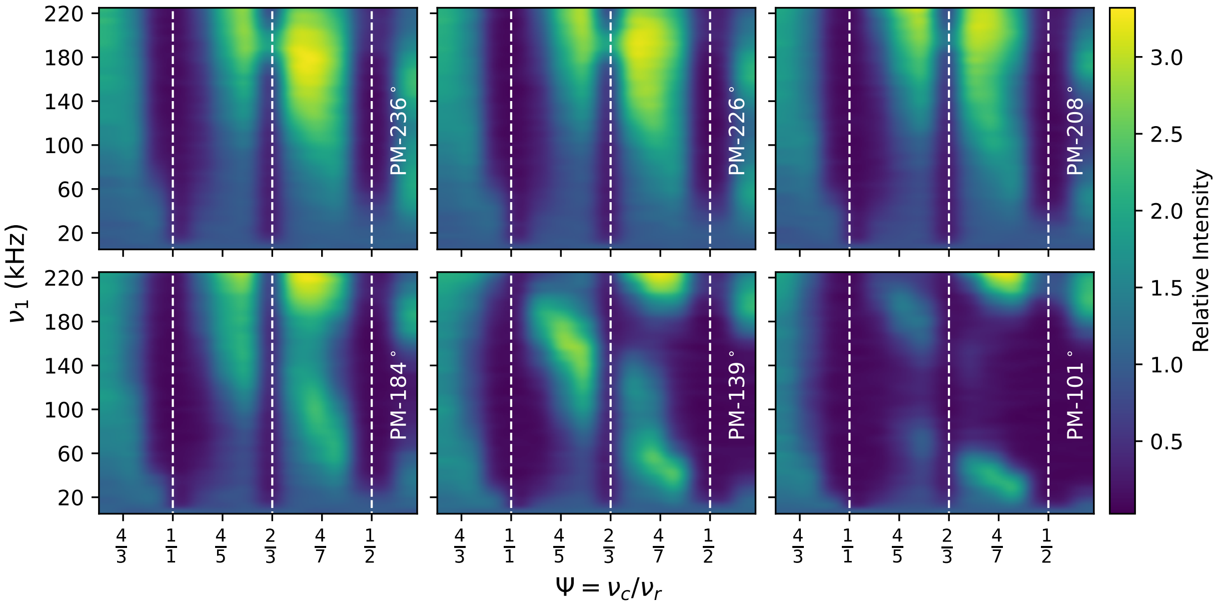


FIG. S3. Experimental data for the optimization of homonuclear decoupling at the MAS frequency of 95.24 kHz on a sample of the tripeptide f-MLF, for the homonuclear decoupling sequence PM-$\theta$, as indicated for each subplot (see main text for the nomenclature). $\Psi = \nu_c/\nu_r$ and the rf-amplitude are independently varied. The color-scale represents the intensity of the resonance observed with homonuclear decoupling after a 3.0 ms echo, normalized by the intensity in the absence of homonuclear decoupling for the same echo duration, for the leucine residue in f-MLF. A windowless version of PM-$\theta$ with a 3-pulse phase ramp is used. Theoretically predicted resonance conditions are indicated by dashed vertical lines.

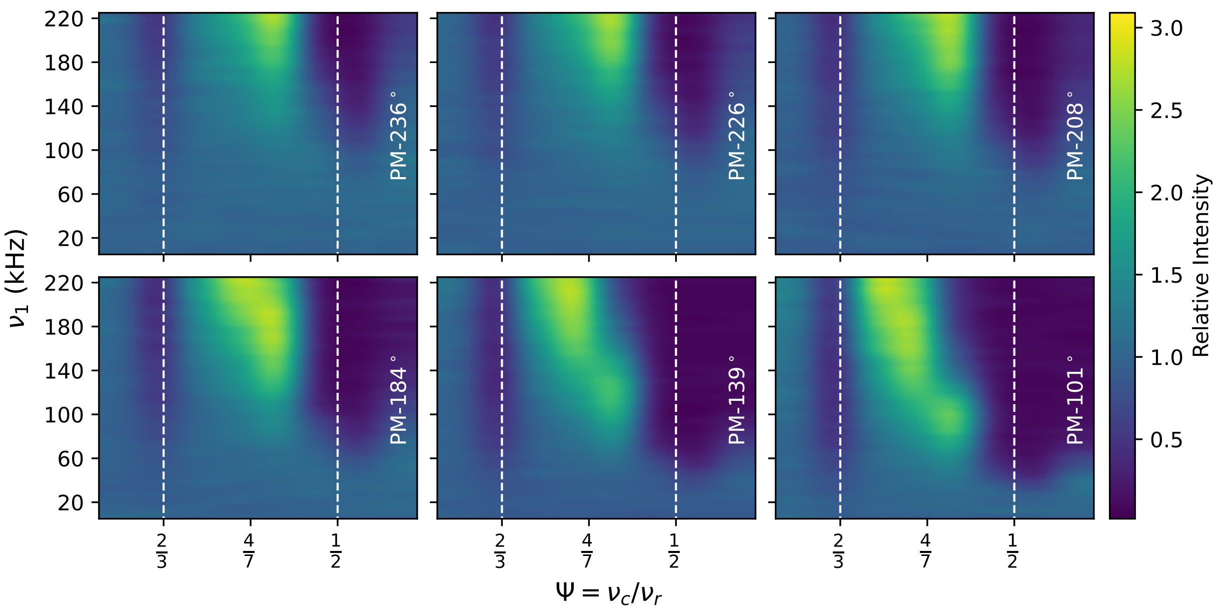


FIG. S4. Experimental data for the optimization of homonuclear decoupling at the MAS frequency of 95.24 kHz on a sample of the tripeptide f-MLF, for the homonuclear decoupling sequence PM-$\theta$, as indicated for each subplot (see main text for the nomenclature). $\Psi = \nu_c/\nu_r$ and the rf-amplitude are independently varied. The color-scale represents the intensity of the resonance observed with homonuclear decoupling after a 3.0 ms echo, normalized by the intensity in the absence of homonuclear decoupling for the same echo duration, for the leucine residue in f-MLF. A windowed version of PM-$\theta$ with two 5.3 $\mu$s windows per supercycle and a 3-pulse phase ramp is used for these plots. Theoretically predicted resonance conditions are indicated by dashed vertical lines.

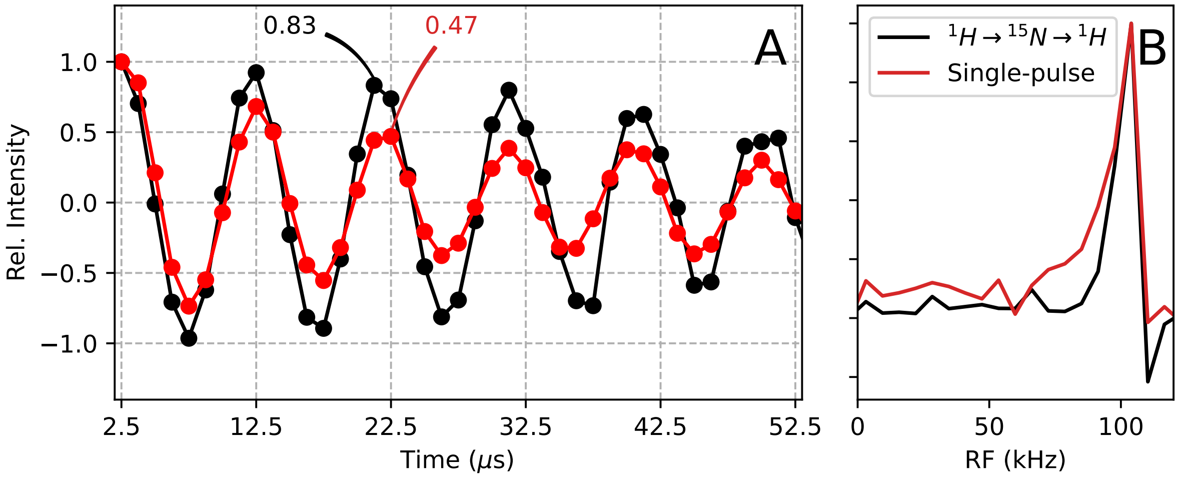


FIG. S5. (A) $^{1}$H nutation profiles for a single pulse experiment at the nominal rf-amplitude of 100 kHz for signal obtained from a $^{15}$N-filtered $^{1}$H experiment (black) and a single pulse $^{1}$H experiment (red). The ratio of the signal intensity for a 810°pulse to that of a 90° pulse, often taken as an indicator of the degree of rf-homogeneity, are 83% and 47% respectively. (B) Fourier transform of the nutation profiles in (A) shows a larger degree of rf-inhomogeneity in a single pulse experiment.

### S3. COHERENCE LIFETIMES UNDER HOMONUCLEAR DECOUPLING

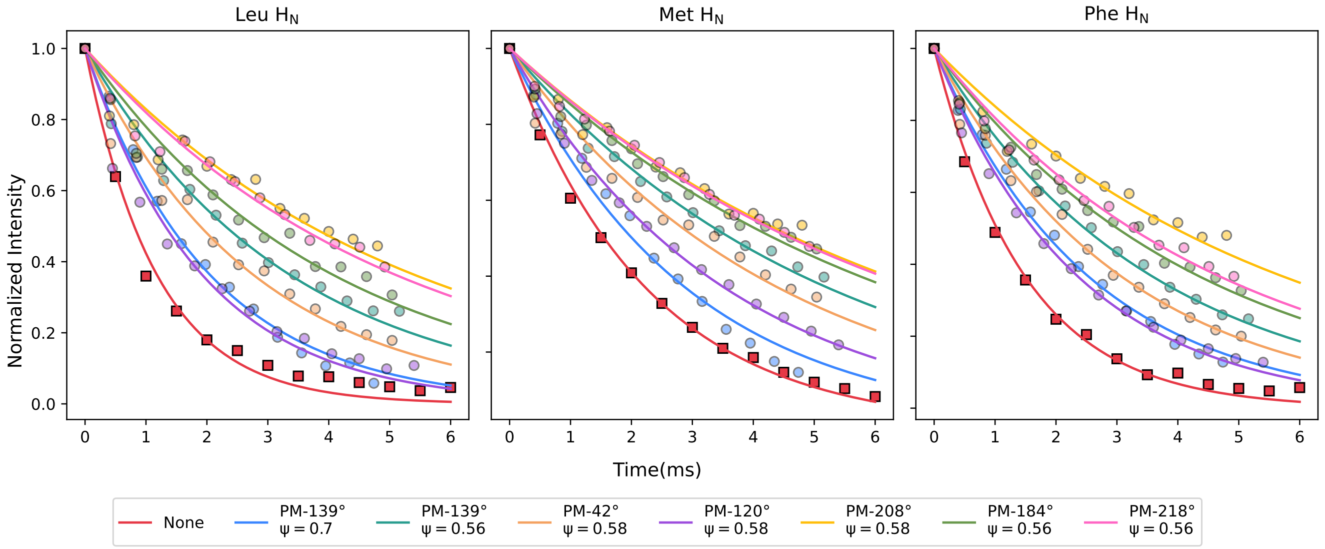


FIG. S6. Signal intensity as a function of spin-echo duration for the three amide protons of f-MLF at a MAS frequency of 62.50 kHz. Supercycled windowed PM-$\theta$ homonuclear decoupling, with values of $\theta$ and $\Psi$ indicated in the legend, was applied during the echo period. Two windows of 5.3 $\mu$s each were included per supercycle. The time axis for each dataset is scaled by the scaling factor ($\lambda$) so that the decays reflect effective improvements in coherence times. Data without homonuclear decoupling are represented by squares. All experimental conditions, $T_2$' times, and scaling factors are tabulated below.

TABLE S2. Experimental conditions for homonuclear decoupling, $T_2'$ times, and scaling factors for f-MLF amide proton resonances, corresponding to data shown in Figure S6.

| Peak | RF (kHz) | $\Psi$ | $\theta$ (°) | $T_2'$ (ms) | $\lambda$ | $T_2' \cdot \lambda$ (ms) |
|---|---|---|---|---|---|---|
| M1 | 0 | – | – | 2.2 | 1.00 | 2.2 |
| L2 | 0 | – | – | 1.2 | 1.00 | 1.2 |
| F3 | 0 | – | – | 1.5 | 1.00 | 1.5 |
| M1 | 45 | 0.58 | 42 | 5.3 | 0.84 | 4.5 |
| L2 | 45 | 0.58 | 42 | 3.2 | 0.84 | 2.7 |
| F3 | 45 | 0.58 | 42 | 3.6 | 0.84 | 3.0 |
| M1 | 45 | 0.58 | 120 | 3.9 | 0.90 | 3.5 |
| L2 | 45 | 0.58 | 120 | 2.1 | 0.90 | 1.9 |
| F3 | 45 | 0.58 | 120 | 2.6 | 0.90 | 2.3 |
| M1 | 50 | 0.56 | 139 | 6.1 | 0.86 | 5.2 |
| L2 | 50 | 0.56 | 139 | 3.9 | 0.86 | 3.4 |
| F3 | 50 | 0.56 | 139 | 4.2 | 0.86 | 3.6 |
| M1 | 70 | 0.56 | 184 | 7.5 | 0.84 | 6.3 |
| L2 | 70 | 0.56 | 184 | 4.8 | 0.84 | 4.0 |
| F3 | 70 | 0.56 | 184 | 5.2 | 0.84 | 4.4 |
| M1 | 100 | 0.56 | 218 | 8.1 | 0.82 | 6.6 |
| L2 | 100 | 0.56 | 218 | 6.1 | 0.82 | 5.0 |
| F3 | 100 | 0.56 | 218 | 5.7 | 0.82 | 4.7 |
| M1 | 120 | 0.70 | 139 | 3.7 | 0.79 | 2.9 |
| L2 | 120 | 0.70 | 139 | 2.6 | 0.79 | 2.1 |
| F3 | 120 | 0.70 | 139 | 3.2 | 0.79 | 2.5 |
| M1 | 120 | 0.58 | 208 | 8.5 | 0.80 | 6.8 |
| L2 | 120 | 0.58 | 208 | 6.7 | 0.80 | 5.4 |
| F3 | 120 | 0.58 | 208 | 7.1 | 0.80 | 5.7 |

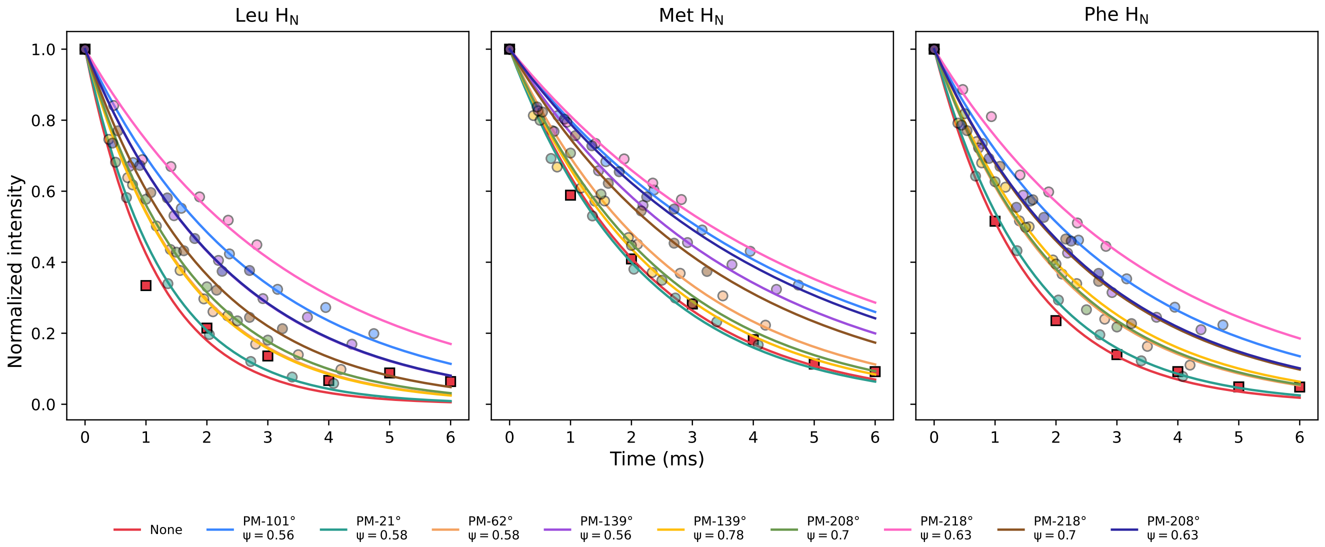


FIG. S7. Signal intensity as a function of spin-echo duration for the three amide protons of f-MLF at a MAS frequency of 62.50 kHz. Supercycled non-windowed PM-$\theta$ homonuclear decoupling, with values of $\theta$ and $\Psi$ indicated in the legend, was applied during the echo period. The time axis for each dataset is scaled by the scaling factor ($\lambda$) so that the decays reflect effective improvements in coherence times. Data without homonuclear decoupling are represented by squares. All experimental conditions, $T_2$' times, and scaling factors are tabulated below.

TABLE S3. Experimental conditions for homonuclear decoupling, $T_2'$ times, and scaling factors for f-MLF amide proton resonances, corresponding to data shown in Figure S7.

| Peak | RF (kHz) | $\Psi$ | $\theta$ (°) | $T_2'$ (ms) | $\lambda$ | $T_2' \cdot \lambda$ (ms) |
|---|---|---|---|---|---|---|
| M1 | 0 | - | - | 2.2 | 1 | 2.2 |
| L2 | 0 | - | - | 1.2 | 1 | 1.2 |
| F3 | 0 | - | - | 1.5 | 1 | 1.5 |
| M1 | 25 | 0.56 | 101 | 5.6 | 0.79 | 4.4 |
| L2 | 25 | 0.56 | 101 | 3.3 | 0.79 | 2.6 |
| F3 | 25 | 0.56 | 101 | 3.8 | 0.79 | 3.0 |
| M1 | 30 | 0.58 | 21 | 3.1 | 0.68 | 2.1 |
| L2 | 30 | 0.58 | 21 | 1.8 | 0.68 | 1.2 |
| F3 | 30 | 0.58 | 21 | 2.4 | 0.68 | 1.6 |
| M1 | 30 | 0.58 | 62 | 3.9 | 0.7 | 2.7 |
| L2 | 30 | 0.58 | 62 | 2.3 | 0.7 | 1.6 |
| F3 | 30 | 0.58 | 62 | 3 | 0.7 | 2.1 |
| M1 | 35 | 0.56 | 139 | 5.1 | 0.73 | 3.7 |
| L2 | 35 | 0.56 | 139 | 3 | 0.73 | 2.2 |
| F3 | 35 | 0.56 | 139 | 3.5 | 0.73 | 2.6 |
| M1 | 90 | 0.78 | 139 | 6.2 | 0.39 | 2.4 |
| L2 | 90 | 0.78 | 139 | 4.2 | 0.39 | 1.6 |
| F3 | 90 | 0.78 | 139 | 5.6 | 0.39 | 2.2 |
| M1 | 130.8 | 0.7 | 208 | 5.1 | 0.5 | 2.6 |
| L2 | 130.8 | 0.7 | 208 | 3.5 | 0.5 | 1.8 |
| F3 | 130.8 | 0.7 | 208 | 4.2 | 0.5 | 2.1 |
| M1 | 130.8 | 0.63 | 218 | 10.2 | 0.47 | 4.8 |
| L2 | 130.8 | 0.63 | 218 | 7.2 | 0.47 | 3.4 |
| F3 | 130.8 | 0.63 | 218 | 7.6 | 0.47 | 3.6 |
| M1 | 130.8 | 0.7 | 218 | 6.4 | 0.54 | 3.5 |
| L2 | 130.8 | 0.7 | 218 | 3.7 | 0.54 | 2.0 |
| F3 | 130.8 | 0.7 | 218 | 4.8 | 0.54 | 2.6 |
| M1 | 131.3 | 0.63 | 208 | 9.4 | 0.45 | 4.2 |
| L2 | 131.3 | 0.63 | 208 | 5.3 | 0.45 | 2.4 |
| F3 | 131.3 | 0.63 | 208 | 5.8 | 0.45 | 2.6 |

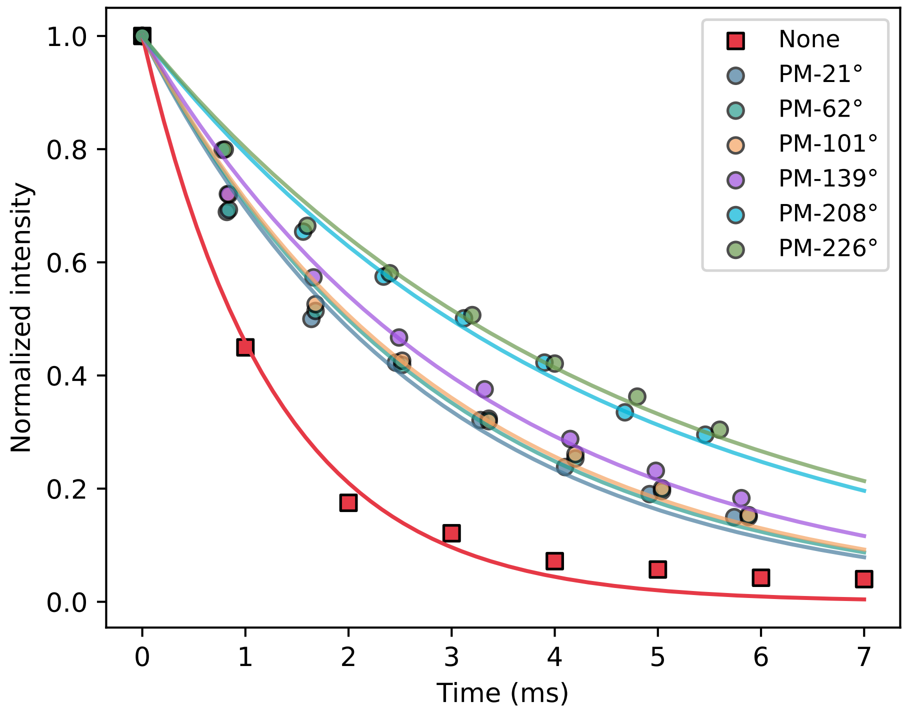


FIG. S8. Signal intensity as a function of spin-echo duration for the amide protons of GB1 at a MAS frequency of 62.50 kHz. Supercycled windowed PM-$\theta$ homonuclear decoupling, with values of $\theta$ indicated in the legend, was applied during the echo period. Two windows of 5.3 $\mu$s each were included per supercycle. The time axis for each dataset is scaled by the scaling factor ($\lambda$) so that the decays reflect the effective improvement in coherence times. Data without homonuclear decoupling are represented by squares. All experimental conditions, $T_2$' times, and scaling factors are tabulated below.

TABLE S4. Experimental conditions for homonuclear decoupling, $T_2'$ times, and scaling factors for the bulk amide proton resonances in GB1, corresponding to data shown in Figure S8.

| RF (kHz) | $\Psi$ | $\theta$ (°) | $T_2'$ (ms) | $\lambda$ | $T_2' \cdot \lambda$ (ms) |
|---|---|---|---|---|---|
| 0 | – | – | 1.3 | 1.00 | 1.3 |
| 40 | 0.58 | 21 | 3.4 | 0.82 | 2.8 |
| 40 | 0.58 | 62 | 3.4 | 0.84 | 2.9 |
| 45 | 0.58 | 101 | 3.5 | 0.84 | 2.9 |
| 55 | 0.57 | 139 | 3.9 | 0.83 | 3.2 |
| 120 | 0.58 | 208 | 5.5 | 0.78 | 4.3 |
| 125 | 0.58 | 226 | 5.7 | 0.80 | 4.6 |

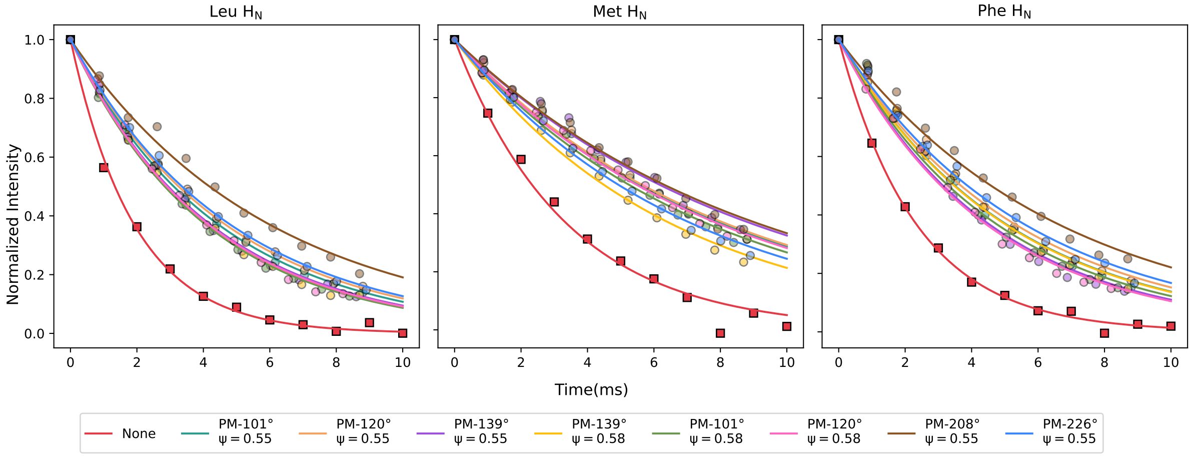


FIG. S9. Signal intensity as a function of spin-echo duration for the three amide protons of f-MLF at a MAS frequency of 95.24 kHz. Supercycled windowed PM-$\theta$ homonuclear decoupling, with values of $\theta$ indicated in the legend, was applied during the echo period. Two windows of 5.3 $\mu$s each were included per supercycle. The time axis for each dataset is scaled by the scaling factor ($\lambda$) so that the decays reflect the effective improvement in coherence times. Data without homonuclear decoupling are represented by squares. All experimental conditions, $T_2$' times, and scaling factors are tabulated below.

TABLE S5. Experimental conditions for homonuclear decoupling, $T_2'$ times, and scaling factors for f-MLF amide proton resonances, corresponding to data shown in Figure S9.

| Peak | RF (kHz) | $\Psi$ | $\theta$ (°) | $T_2'$ (ms) | $\lambda$ | $T_2' \cdot \lambda$ (ms) |
|---|---|---|---|---|---|---|
| L2 | 0 | - | - | 2.0 | 1 | 2.0 |
| M1 | 0 | - | - | 3.7 | 1 | 3.7 |
| F3 | 0 | - | - | 2.3 | 1 | 2.3 |
| L2 | 109 | 0.55 | 101 | 5.1 | 0.88 | 4.5 |
| M1 | 109 | 0.55 | 101 | 9.1 | 0.88 | 8.0 |
| F3 | 109 | 0.55 | 101 | 5.7 | 0.88 | 5.0 |
| L2 | 117 | 0.55 | 120 | 5.3 | 0.88 | 4.7 |
| M1 | 117 | 0.55 | 120 | 9.2 | 0.88 | 8.1 |
| F3 | 117 | 0.55 | 120 | 6.0 | 0.88 | 5.3 |
| L2 | 149 | 0.55 | 139 | 4.9 | 0.86 | 4.2 |
| M1 | 149 | 0.55 | 139 | 10.3 | 0.86 | 8.9 |
| F3 | 149 | 0.55 | 139 | 5.3 | 0.86 | 4.6 |
| L2 | 170 | 0.58 | 139 | 4.7 | 0.87 | 4.1 |
| M1 | 170 | 0.58 | 139 | 7.4 | 0.87 | 6.4 |
| F3 | 170 | 0.58 | 139 | 5.8 | 0.87 | 5.1 |
| L2 | 175 | 0.58 | 101 | 4.9 | 0.84 | 4.1 |
| M1 | 175 | 0.58 | 101 | 9.1 | 0.84 | 7.6 |
| F3 | 175 | 0.58 | 101 | 5.7 | 0.84 | 4.8 |
| L2 | 203 | 0.58 | 120 | 5.1 | 0.82 | 4.2 |
| M1 | 203 | 0.58 | 120 | 9.6 | 0.82 | 7.9 |
| F3 | 203 | 0.58 | 120 | 5.4 | 0.82 | 4.4 |
| L2 | 219 | 0.55 | 208 | 6.9 | 0.87 | 6.0 |
| M1 | 219 | 0.55 | 208 | 10.5 | 0.87 | 9.1 |
| F3 | 219 | 0.55 | 208 | 7.6 | 0.87 | 6.6 |
| L2 | 219 | 0.55 | 226 | 5.4 | 0.89 | 4.8 |
| M1 | 219 | 0.55 | 226 | 8.0 | 0.89 | 7.1 |
| F3 | 219 | 0.55 | 226 | 6.3 | 0.89 | 5.6 |

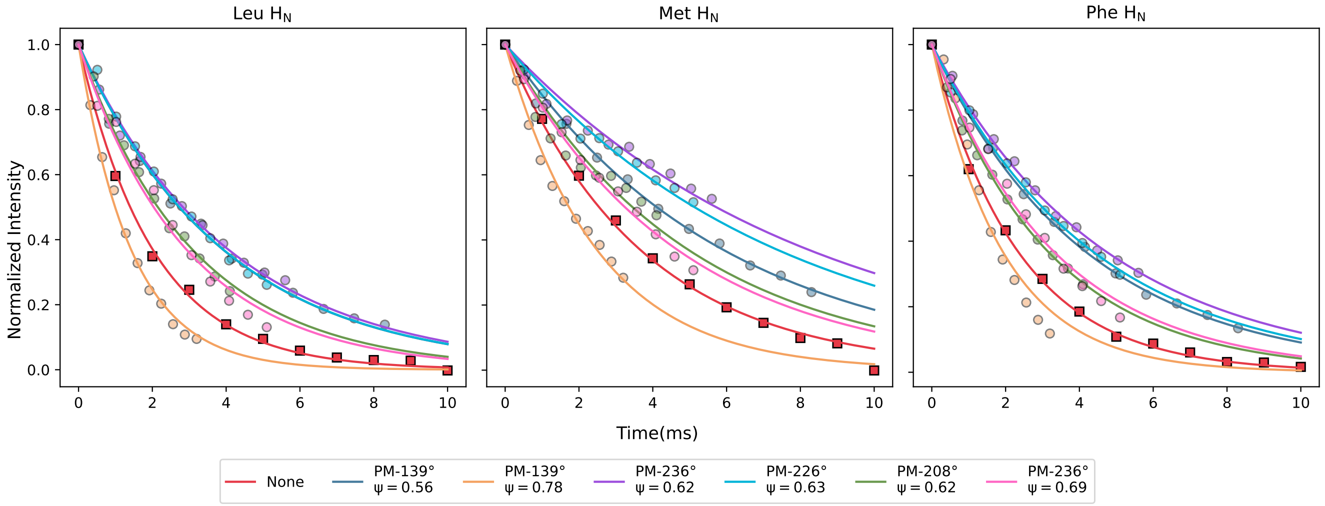


FIG. S10. Signal intensity as a function of spin-echo duration for the three amide protons of f-MLF at a MAS frequency of 95.24 kHz. Supercycled non-windowed PM-$\theta$ homonuclear decoupling, with values of $\theta$ indicated in the legend, was applied during the echo period. The time axis for each dataset is scaled by the scaling factor ($\lambda$) so that the decays reflect the effective improvement in coherence times. Data without homonuclear decoupling are represented by squares. All experimental conditions, $T_2$' times, and scaling factors are tabulated below. Note that in spite of the long apparent $T_2$' observed for one of the conditions (PM-139°at $\Psi$=0.78), there is a decrease in coherence time due to a low scaling factor (see tabulated values below).

TABLE S6. Experimental conditions for homonuclear decoupling, $T_2'$ times, and scaling factors for f-MLF amide proton resonances, corresponding to data shown in Figure S10.

| Peak | RF (kHz) | $\Psi$ | $\theta$ (°) | $T_2'$ (ms) | $\lambda$ | $T_2' \cdot \lambda$ (ms) |
|---|---|---|---|---|---|---|
| M1 | 0 | - | - | 3.7 | 1 | 3.7 |
| L2 | 0 | - | - | 2.0 | 1 | 2.0 |
| F3 | 0 | - | - | 2.3 | 1 | 2.3 |
| M1 | 45 | 0.56 | 139 | 7.2 | 0.83 | 6.0 |
| L2 | 45 | 0.56 | 139 | 4.7 | 0.83 | 3.9 |
| F3 | 45 | 0.56 | 139 | 5.0 | 0.83 | 4.2 |
| M1 | 170 | 0.78 | 139 | 7.8 | 0.32 | 2.5 |
| L2 | 170 | 0.78 | 139 | 4.5 | 0.32 | 1.4 |
| F3 | 170 | 0.78 | 139 | 6.0 | 0.32 | 1.9 |
| M1 | 170 | 0.62 | 236 | 14.8 | 0.56 | 8.3 |
| L2 | 170 | 0.62 | 236 | 7.3 | 0.56 | 4.1 |
| F3 | 170 | 0.62 | 236 | 8.4 | 0.56 | 4.7 |
| M1 | 190 | 0.63 | 226 | 14.5 | 0.51 | 7.4 |
| L2 | 190 | 0.63 | 226 | 7.7 | 0.51 | 3.9 |
| F3 | 190 | 0.63 | 226 | 8.6 | 0.51 | 4.4 |
| M1 | 208 | 0.62 | 208 | 12.2 | 0.41 | 5 |
| L2 | 208 | 0.62 | 208 | 7.6 | 0.41 | 3.1 |
| F3 | 208 | 0.62 | 208 | 7.7 | 0.41 | 3.2 |
| M1 | 211 | 0.69 | 236 | 9.2 | 0.51 | 4.7 |
| L2 | 211 | 0.69 | 236 | 5.8 | 0.51 | 3.0 |
| F3 | 211 | 0.69 | 236 | 6.5 | 0.51 | 3.3 |

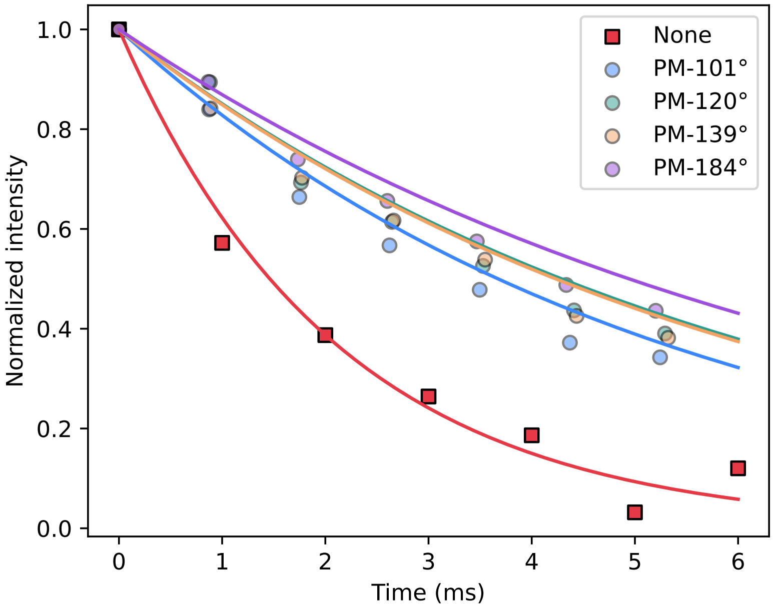


FIG. S11. Signal intensity as a function of spin-echo duration for the amide protons of GB1 at a MAS frequency of 95.24 kHz. Supercycled windowed PM-$\theta$ homonuclear decoupling, with values of $\theta$ indicated in the legend, was applied during the echo period. Two windows of 5.3 $\mu$s each were included per supercycle. The time axis for each dataset is scaled by the scaling factor ($\lambda$) so that the decays reflect the effective improvement in coherence times. Data without homonuclear decoupling are represented by squares. All experimental conditions, $T_2$' times, and scaling factors are tabulated below.

TABLE S7. Experimental conditions for homonuclear decoupling, $T_2'$ times, and scaling factors for bulk amide proton signal in GB1, corresponding to data shown in Figure S11.

| RF (kHz) | $\Psi$ | $\theta$ (°) | $T_2'$ (ms) | $\lambda$ | $T_2' \cdot \lambda$ (ms) |
|---|---|---|---|---|---|
| 0 | - | - | 2.1 | 1 | 2.1 |
| 116 | 0.55 | 101 | 5.3 | 0.87 | 4.6 |
| 124 | 0.55 | 120 | 6.2 | 0.88 | 5.5 |
| 138 | 0.55 | 139 | 6.1 | 0.89 | 5.4 |
| 196 | 0.55 | 184 | 7.1 | 0.87 | 6.2 |

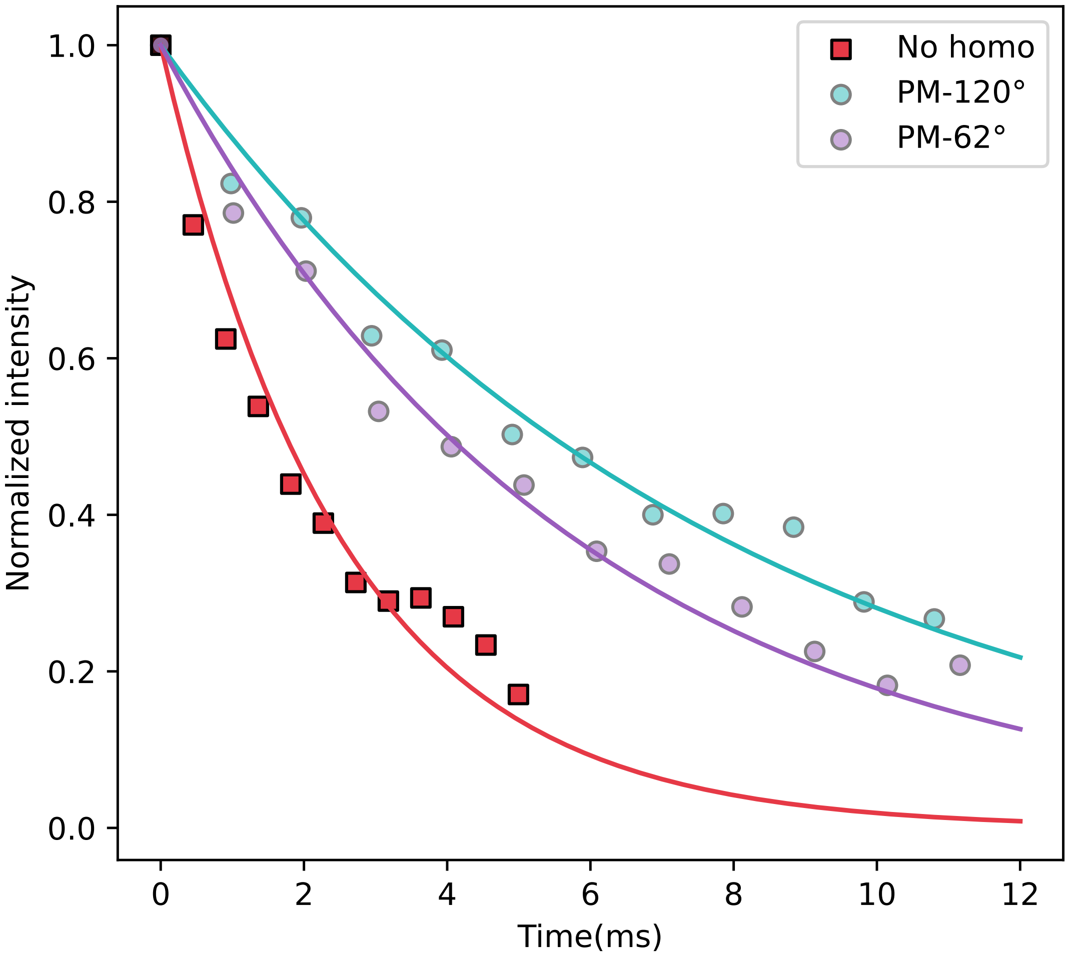


FIG. S12. Signal intensity as a function of spin-echo duration for the amide protons of ParM at a MAS frequency of 95.24 kHz. Supercycled windowed PM-$\theta$ homonuclear decoupling, with values of $\theta$ indicated in the legend, was applied during the echo period. Two windows of 5.3 $\mu$s each were included per supercycle. The time axis for each dataset is scaled by the scaling factor ($\lambda$) so that the decays reflect the effective improvement in coherence times. Data without homonuclear decoupling are represented by squares. All experimental conditions, $T_2$' times, and scaling factors are tabulated below.

TABLE S8. Experimental conditions for homonuclear decoupling, $T_2'$ times, and scaling factors for bulk amide proton signal in ParM, corresponding to data shown in Figure S12.

| RF (kHz) | $\Psi$ | $\theta$ (°) | $T_2'$ (ms) | $\lambda$ | $T_2' \cdot \lambda$ (ms) |
|---|---|---|---|---|---|
| 0 | - | - | 2.5 | 1 | 2.5 |
| 82 | 0.55 | 62 | 6.4 | 0.93 | 6 |
| 123 | 0.55 | 120 | 8.8 | 0.9 | 7.9 |

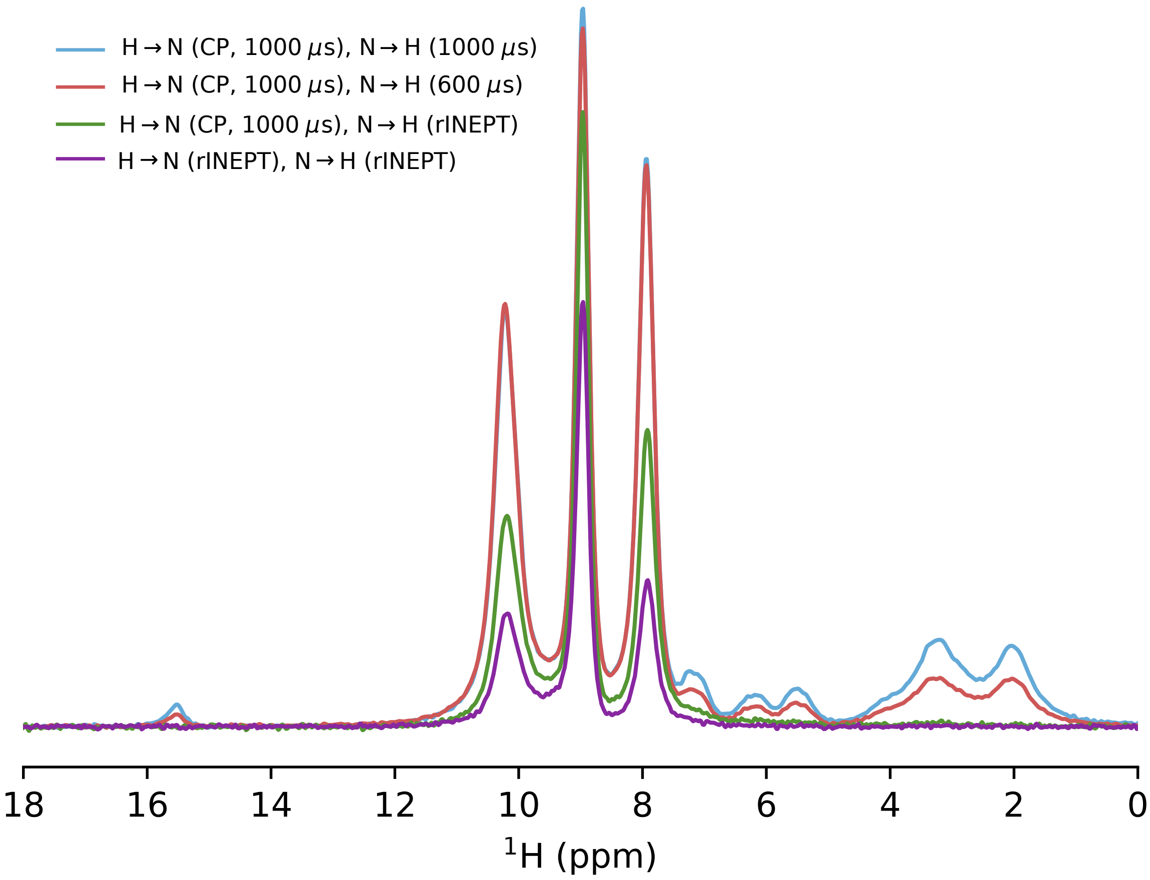


FIG. S13. Comparison of efficiency of INEPT transfer in the presence of homonuclear decoupling with that obtained in a CP-based experiment for amide protons of f-MLF at 95.24 kHz MAS. Non-windowed homonuclear decoupling sequence used here is PM-236°($\Psi$ = 0.62, rf = 170 kHz, $\lambda$ = 0.56), and the effective $T_2$' values for the three peaks are 4.1, 8.3, and 4.7 ms respectively. For the CP-based experiment, forward and reverse CP contact times were both kept at 1.0 ms and 1.0 ms (blue), or 1.0 ms and 0.6 ms (red), respectively. These were optimized for the highest sensitivity of the amide peaks rather than optimal selectivity for the transfer. The effective 1/4J transfer times during rINEPT were optimized to 2.25 ms when the transverse magnetization is on $^1$H and 3.25 ms when it is on $^{15}$N. Note that longer times are required due to the lower scaling factor of this sequence, which scales both chemical shifts and scalar coupling identically. Spectrum in purple was obtained with both the transfers based on rINEPT and the one in green was obtained with only the transfer from $^{15}$N to $^1$H implemented using rINEPT. All the spectra were obtained using 1024 scans and all other relevant experimental parameters ($^{15}$N decoupling, acquisition times, recycle delays) kept identical. The net efficiency with respect to CP based transfers for transfers based on only rINEPT or a combination of CP and rINEPT were as follows: L2: 27%, 55%, M1: 60%, 84% F3: 25%, 50%.

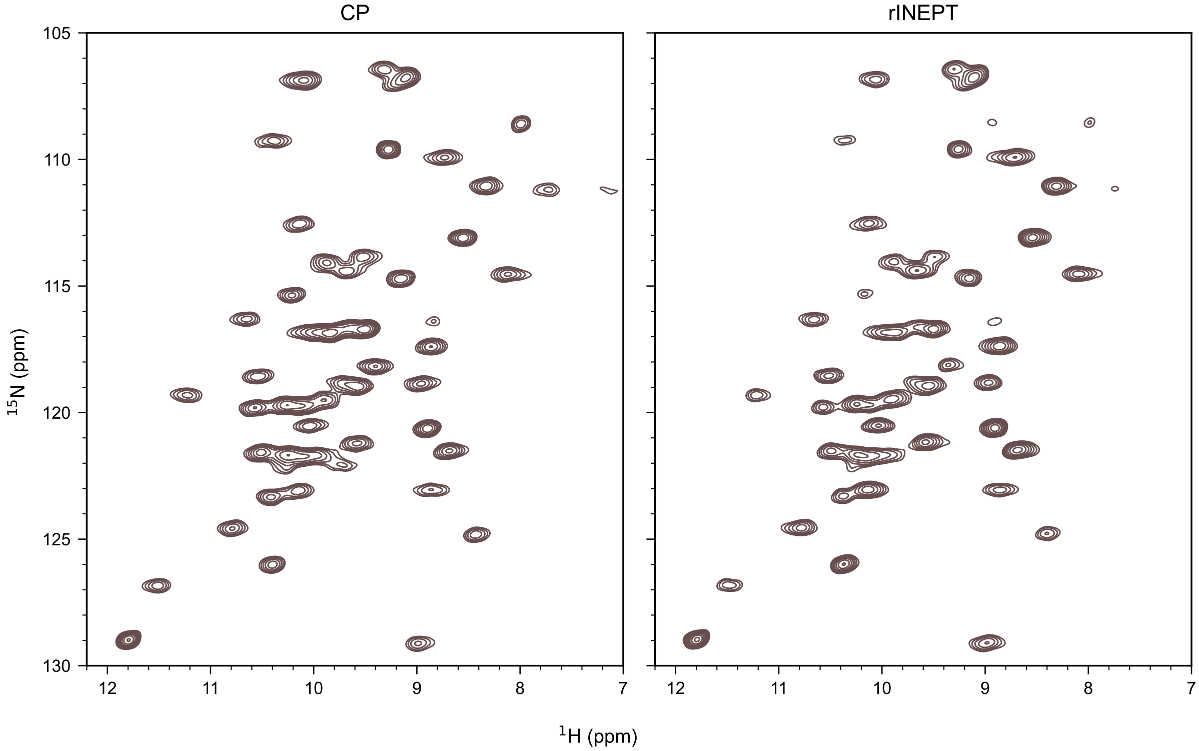


FIG. S14. Comparison of 2D $^{15}$N-$^{1}$H spectra of GB1 obtaiend using either CP-based transfers or rINEPT-based transfer (in the presence of homonuclear decoupling) at the spinning frequency of 95.24 kHz. Non-windowed homonuclear decoupling sequence similar to that used for f-MLF (Figure S13) is used for this experiment (PM-236°, $\Psi$ = 0.62, rf = 170 kHz, $\lambda$ = 0.56), The sensitivity of the rINEPT experiment as compared to the CP-based experiment was 25%. The above spectra were obtained with 8 scans (CP) and 128 scans (rINEPT) to compensate for this difference. Spectra were processed identically with window functions in both the direct and indirect dimensions, and are plotted after dividing by the rmsd of the noise and $\sqrt{ns}$, where $ns$ is the number of scans. Contours in both the spectra start at the same level and are spaced identically.

### S4. RESOLUTION ENHANCEMENT USING HOMONUCLEAR DECOUPLING AND WINDOWED DETECTION

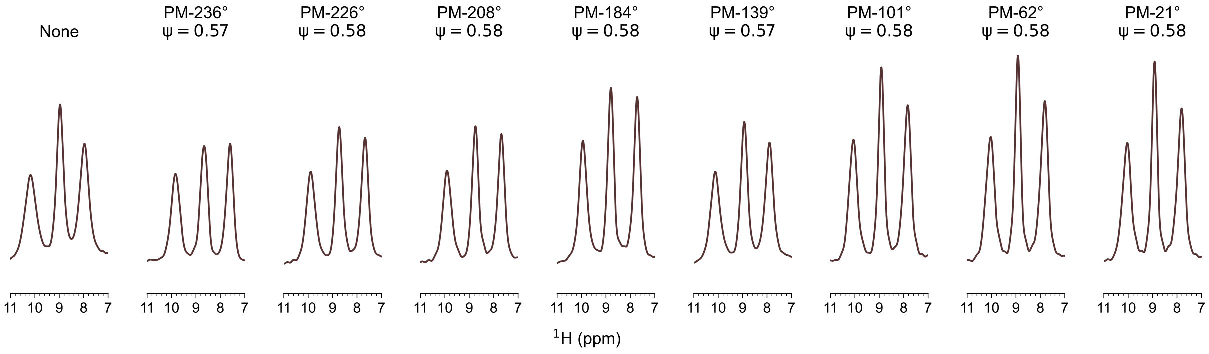


FIG. S15. $^{15}$N edited $^{1}$H 1D spectra of f-MLF recorded at 62.50 kHz MAS on a 1.3 mm HCN using windowed homonuclear decoupling, for the indicated values of the PM-$\theta$ sequences. The control spectrum without homonuclear decoupling is also obtained using windowed sequence to facilitate a direct comparison. All the spectra with homonuclear decoupling are plotted after scaling by the chemical shift scaling factor ($\lambda$). All spectra were recorded with 256 scans with all parameters kept identical, and are plotted after dividing by the noise rmsd and $\sqrt{ns}$ to facilitate a direct comparison. Experimental parameters for the homonuclear decoupling, linewidths for each peak, and scaling factors are tabulated below. Apodization is not used.

TABLE S9. Linewidths of f-MLF amide proton resonances measured without homonuclear decoupling and with windowed homonuclear decoupling under different phase-ramp conditions. The scaling factor is already included in the linewidth, but is reported separately for completeness.

| Peak | RF (kHz) | $\Psi$ | $\theta$ (°) | FWHM (Hz) | $\lambda$ |
|---|---|---|---|---|---|
| L2 | 0 | - | - | 406 | 1 |
| M1 | 0 | - | - | 251 | 1 |
| F3 | 0 | - | - | 293 | 1 |
| L2 | 126 | 0.57 | 236 | 363 | 0.83 |
| M1 | 126 | 0.57 | 236 | 313 | 0.83 |
| F3 | 126 | 0.57 | 236 | 292 | 0.83 |
| L2 | 118 | 0.58 | 226 | 340 | 0.85 |
| M1 | 118 | 0.58 | 226 | 242 | 0.85 |
| F3 | 118 | 0.58 | 226 | 244 | 0.85 |
| L2 | 113 | 0.58 | 208 | 324 | 0.83 |
| M1 | 113 | 0.58 | 208 | 229 | 0.83 |
| F3 | 113 | 0.58 | 208 | 236 | 0.83 |
| L2 | 100 | 0.58 | 184 | 278 | 0.81 |
| M1 | 100 | 0.58 | 184 | 218 | 0.81 |
| F3 | 100 | 0.58 | 184 | 221 | 0.81 |
| L2 | 41 | 0.57 | 139 | 352 | 0.92 |
| M1 | 41 | 0.57 | 139 | 244 | 0.92 |
| F3 | 41 | 0.57 | 139 | 271 | 0.92 |
| L2 | 52 | 0.58 | 101 | 310 | 0.83 |
| M1 | 52 | 0.58 | 101 | 208 | 0.83 |
| F3 | 52 | 0.58 | 101 | 259 | 0.83 |
| L2 | 49 | 0.58 | 62 | 315 | 0.81 |
| M1 | 49 | 0.58 | 62 | 200 | 0.81 |
| F3 | 49 | 0.58 | 62 | 261 | 0.81 |
| L2 | 46 | 0.58 | 21 | 311 | 0.81 |
| M1 | 46 | 0.58 | 21 | 197 | 0.81 |
| F3 | 46 | 0.58 | 21 | 260 | 0.81 |

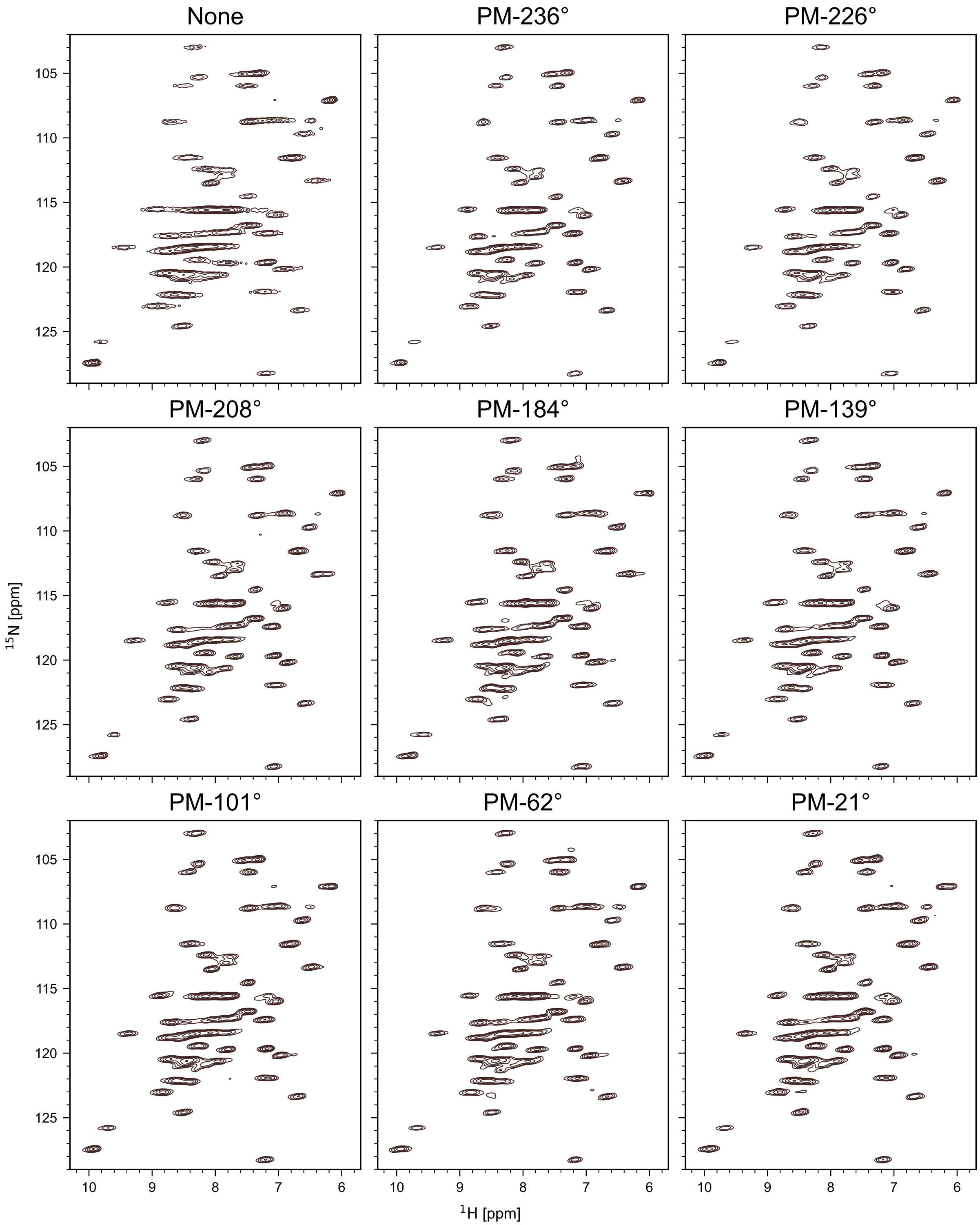


FIG. S16. $^{15}$N edited $^{1}$H 2D spectra of GB1 recorded at 60 kHz MAS on a 1.3 mm HCN probe without homonuclear decoupling (No windowed detection) and with homonuclear decoupling with windowed detection with phase ramps 236°($\psi$ = 0.57, rf = 125 kHz, $\lambda$ = 0.81, NS = 32), 226°($\psi$ = 0.58, rf = 125 kHz, $\lambda$ = 0.82, NS = 32), 208°($\psi$ = 0.58, rf = 120 kHz, $\lambda$ = 0.79, NS = 32), 184°($\psi$ = 0.57, rf = 90 kHz, $\lambda$ = 0.8, NS = 32), 139°($\psi$ = 0.57, rf = 55 kHz, $\lambda$ = 0.83, NS = 32), 101°($\psi$ = 0.58, rf = 45 kHz, $\lambda$ = 0.85, NS = 32), 62°($\psi$ = 0.58, rf = 40 kHz, $\lambda$ = 0.84, NS = 32), 21°($\psi$ = 0.58, rf = 40 kHz, $\lambda$ = 0.82, NS = 32). The spectrum without homonuclear decoupling was recorded with 8 scans(NS). The spectra with homonuclear decoupling are scaled by a chemical shift scaling factor($\lambda$). Spectra are plotted after dividing by the noise rmsd and $\sqrt{ns}$ to facilitate a direct comparison. Apodization is not used in the direct dimension.

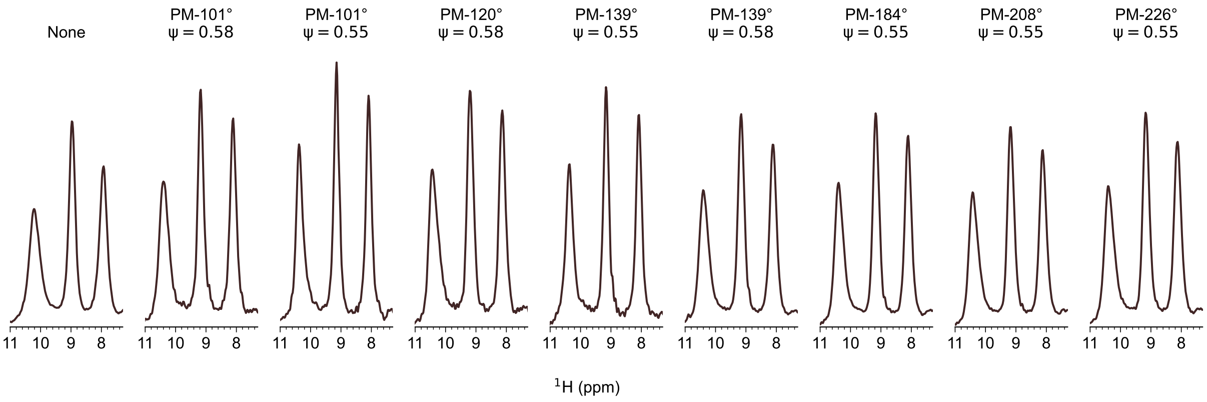


FIG. S17. $^{15}$N edited $^1$H 1D spectra of f-MLF recorded at 95.24 kHz MAS on 0.7 mm HCN probe with homonuclear decoupling and windowed detection. Spectra using the indicated homonuclear decoupling sequence were optimized independently for the best homonuclear and heteronuclear decoupling. All the spectra with homonuclear decoupling are scaled by the chemical shift scaling factor. Spectra are plotted after dividing by the noise rmsd and $\sqrt{ns}$ to facilitate a direct comparison. Note that the spectrum without homonuclear decoupling is also obtained using windowed acquisition sequence to facilitate a clear comparison. Apodization is not used.

TABLE S10. Linewidths of selected f-MLF amide proton resonances measured without homonuclear decoupling and with windowed homonuclear decoupling under different phase-ramp conditions.

| Peak | RF (kHz) | $\Psi$ | $\theta$ (°) | Linewidth (Hz) | Scaling Factor |
|---|---|---|---|---|---|
| L2 | 0 | - | - | 304 | 1 |
| M1 | 0 | - | - | 179 | 1 |
| F3 | 0 | - | - | 204 | 1 |
| L2 | 175 | 0.58 | 101 | 272 | 0.84 |
| M1 | 175 | 0.58 | 101 | 153 | 0.84 |
| F3 | 175 | 0.58 | 101 | 166 | 0.84 |
| L2 | 109 | 0.55 | 101 | 187 | 0.88 |
| M1 | 109 | 0.55 | 101 | 147 | 0.88 |
| F3 | 109 | 0.55 | 101 | 147 | 0.88 |
| L2 | 203 | 0.58 | 120 | 276 | 0.82 |
| M1 | 203 | 0.58 | 120 | 177 | 0.82 |
| F3 | 203 | 0.58 | 120 | 173 | 0.82 |
| L2 | 149 | 0.55 | 139 | 197 | 0.86 |
| M1 | 149 | 0.55 | 139 | 148 | 0.86 |
| F3 | 149 | 0.55 | 139 | 153 | 0.86 |
| L2 | 170 | 0.58 | 139 | 267 | 0.87 |
| M1 | 170 | 0.58 | 139 | 173 | 0.87 |
| F3 | 170 | 0.58 | 139 | 185 | 0.87 |
| L2 | 185 | 0.55 | 184 | 251 | 0.90 |
| M1 | 185 | 0.55 | 184 | 173 | 0.90 |
| F3 | 185 | 0.55 | 184 | 175 | 0.90 |
| L2 | 219 | 0.55 | 208 | 277 | 0.87 |
| M1 | 219 | 0.55 | 208 | 184 | 0.87 |
| F3 | 219 | 0.55 | 208 | 190 | 0.87 |
| L2 | 219 | 0.55 | 226 | 281 | 0.89 |
| M1 | 219 | 0.55 | 226 | 182 | 0.89 |
| F3 | 219 | 0.55 | 226 | 193 | 0.89 |

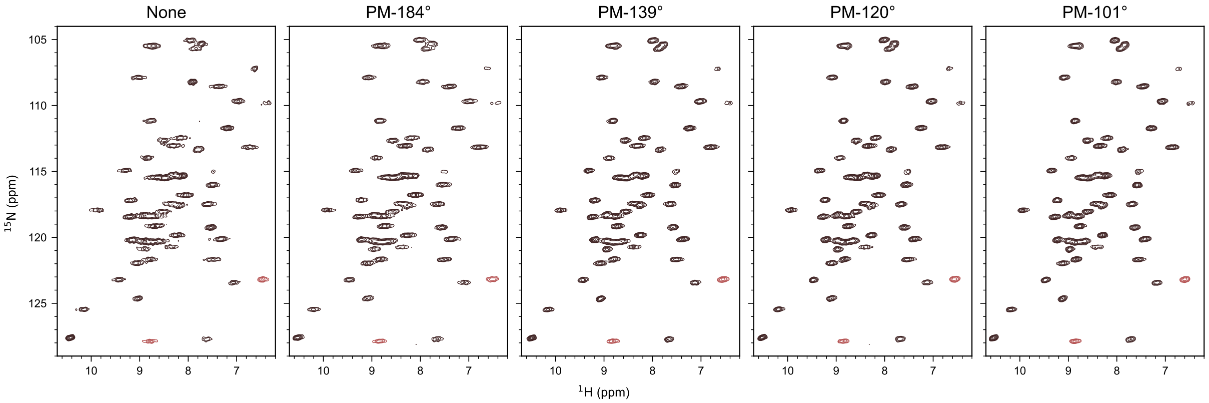


FIG. S18. $^{15}$N edited $^{1}$H 2D spectra of GB1 recorded at 95.24 kHz MAS on a 0.7 mm HCN probe with homonuclear decoupling and windowed detection with the following conditions phase ramps 139°($\psi$ = 0.55, rf = 140 kHz, $\lambda$ = 0.87), 101°($\psi$ = 0.55, rf = 110 kHz, $\lambda$ = 0.83), 62°($\psi$ = 0.55, rf = 100 kHz, $\lambda$ = 0.88) and 21°($\psi$ = 0.55, rf = 90 kHz, $\lambda$ = 0.89). The spectrum without homonuclear decoupling was recorded with 8 scans using normal acquisition, while those with homonuclear decoupling were recorded with 32 scans and windowed acquisition. The spectra with homonuclear decoupling are scaled by a chemical shift scaling factor($\lambda$). Spectra are plotted after dividing by the noise rmsd and $\sqrt{ns}$ to facilitate a direct comparison. Apodization is not used in the direct dimension.

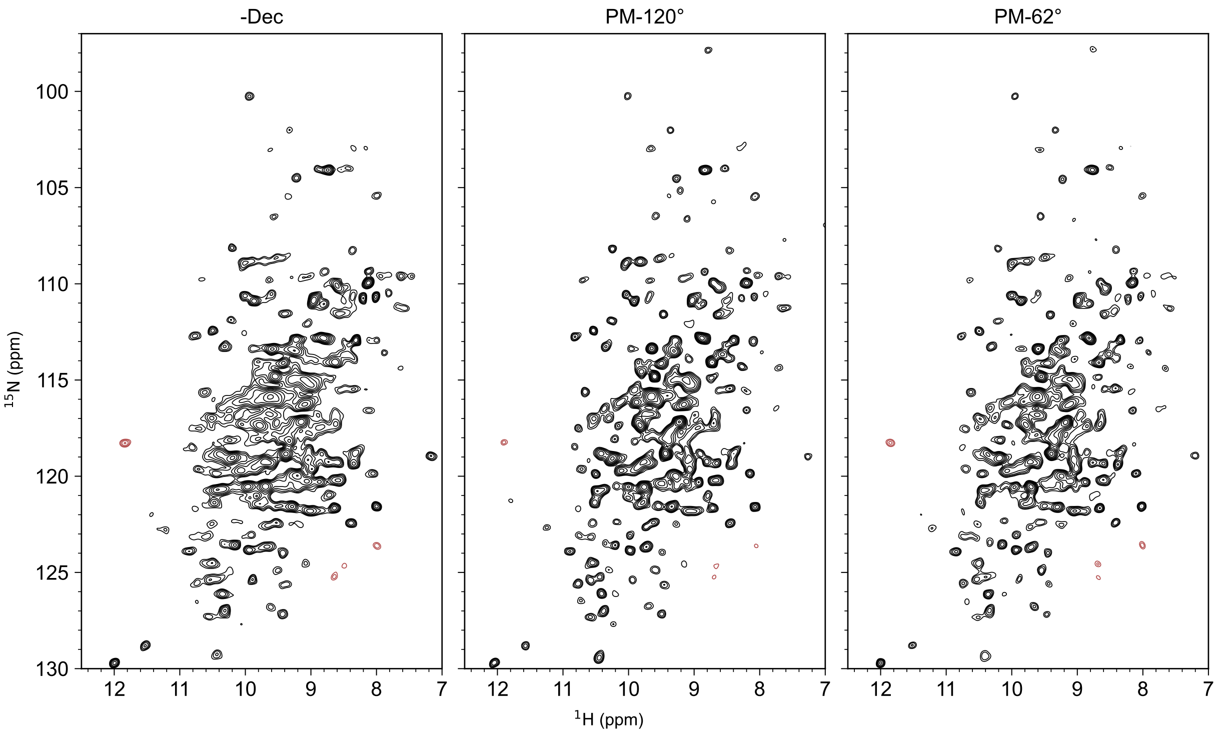


FIG. S19. $^{15}$N edited $^{1}$H 2D spectra of ParM recorded at 95.24 kHz MAS on a 0.7 mm HCN probe with homonuclear decoupling and windowed detection with the following conditions: PM-120°($\Psi$ = 0.55, rf = 124 kHz, $\lambda$ = 0.90) and PM-62°($\Psi$ = 0.55, rf = 82 kHz, $\lambda$ = 0.93). The spectrum without homonuclear decoupling was recorded with 32 scans (standard acquisition), while those with homonuclear decoupling were recorded with 128 scans (windowed acquisition). The spectra with homonuclear decoupling are scaled by a chemical shift scaling factor. Spectra are plotted after dividing by the noise rmsd and $\sqrt{ns}$ to facilitate a direct comparison. Apodization is not used.

### S5. OPTIMIZATION OF HETERONUCLEAR DECOUPLING

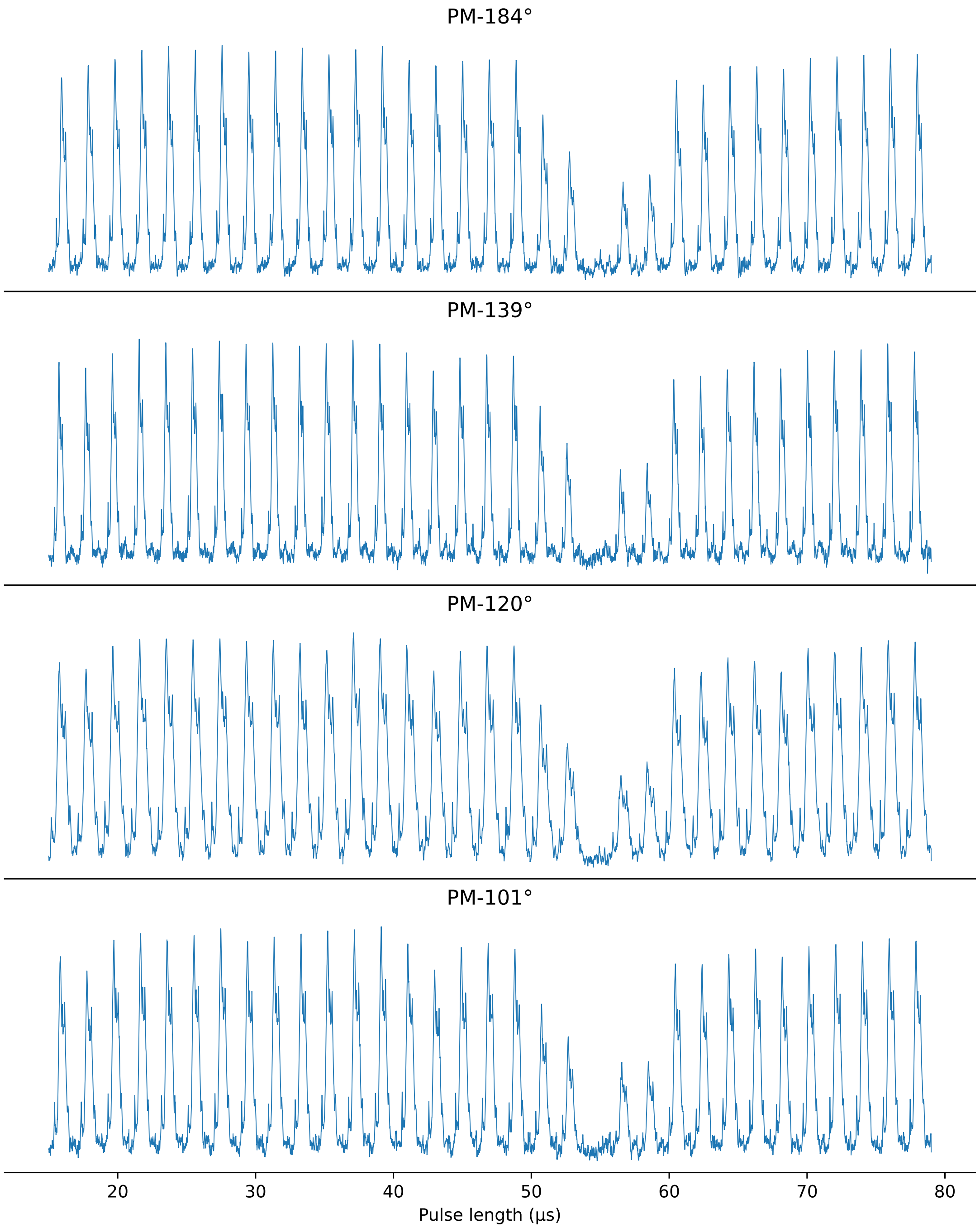


FIG. S20. Parameter optimization of the length of the CW pulse in $rCW^{ApA}$ heteronuclear decoupling. The optimization is shown for $^{15}N$ edited $^{1}H$ 1D spectra of GB1 recorded at 95.2 kHz MAS on a 0.7 mm HCN probe. This optimization is done with homonuclear decoupling pulses during echo time. The echo time is 5 ms. The rf amplitude of $^{15}N$ heteronuclear decoupling was kept 13 kHz and the refocusing pulse in $rCW^{ApA}$ sequence is set to be a $\pi$ pulse in all experiments (38.46 $\mu$s). Optimization is done with different homonuclear decoupling with the following phase ramps: $184°(\psi = 0.55$, rf = 196 kHz), $139°(\psi = 0.55$, rf = 139 kHz), $120°(\psi = 0.55$, rf = 124 kHz), and $101°(\psi = 0.55$, rf = 124 kHz).

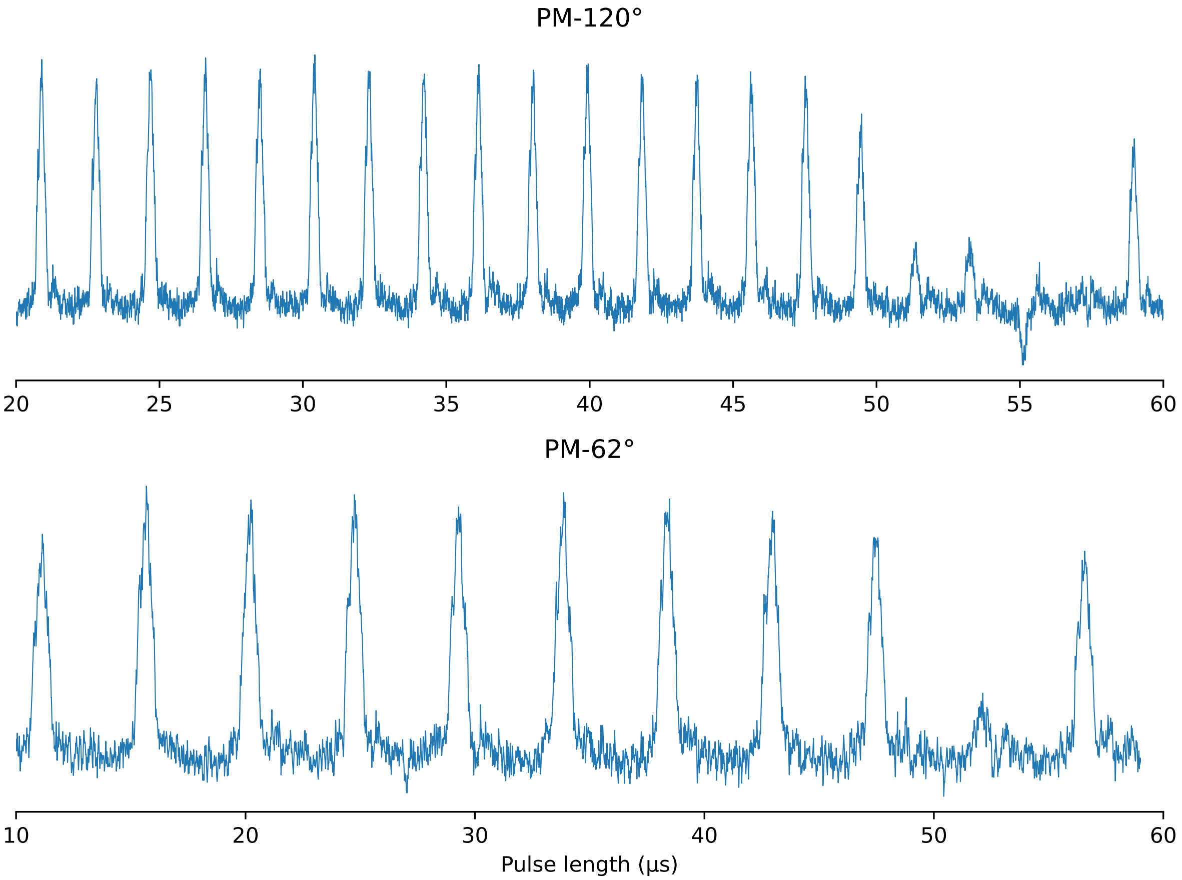


FIG. S21. Parameter optimization of the length of the CW pulse in rCW$^{\text{ApA}}$ heteronuclear decoupling. The optimization is shown for $^{15}$N edited $^{1}$H 1D spectra of ParM recorded at 95.2 kHz MAS on a 0.7 mm HCN probe. This optimization is done with homonuclear decoupling pulses during echo time. The echo time is 5 ms. The rf amplitude of $^{15}$N heteronuclear decoupling was kept 13 kHz and the refocusing pulse in rCW$^{\text{ApA}}$ sequence is set to be a $\pi$ pulse in all experiments (38.46 $\mu$s). Optimization is done with different homonuclear decoupling schemes with the follwoing phase ramps: 120°($\psi = 0.55$, rf = 124 kHz, $\lambda = 0.9$), and 62°($\psi = 0.55$, rf = 82 kHz, $\lambda = 0.932$).

### S6. SENSITIVITY OF SPECTRA ACQUIRED WITH HOMONUCLEAR DECOUPLING AND WINDOWED DETECTION

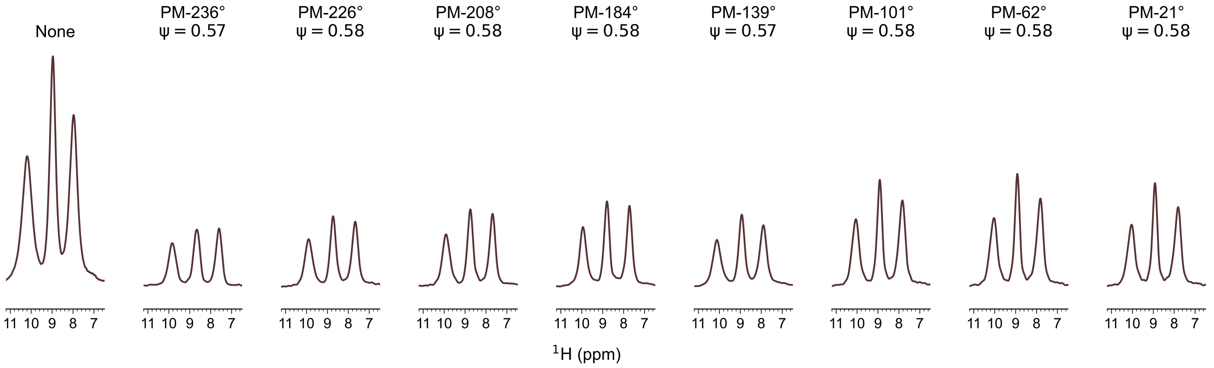


FIG. S22. $^{15}$N edited $^{1}$H 1D spectra of f-MLF recorded at 62.5 kHz MAS on 1.3 mm HCN probe to compare the sensitivity of without homonuclear decoupling (No windowed detection) and with homonuclear decoupling with windowed detection with phase ramps 236°($\psi$ = 0.57, rf = 126 kHz, $\lambda$ = 0.83), 226°($\psi$ = 0.58, rf = 118 kHz, $\lambda$ = 0.85), 208°($\psi$ = 0.58, rf = 113 kHz, $\lambda$ = 0.83), 184°($\psi$ = 0.58, rf = 100 kHz, $\lambda$ = 0.81), 139°($\psi$ = 0.57, rf = 41 kHz, $\lambda$ = 0.92), 101°($\psi$ = 0.58, rf = 52 kHz, $\lambda$ = 0.83), 62°($\psi$ = 0.58, rf = 49 kHz, $\lambda$ = 0.81), and 21°($\psi$ = 0.58, rf = 46 kHz, $\lambda$ = 0.81). All the spectra with homonuclear decoupling are plotted after scaling by the chemical shift scaling factor ($\lambda$). All spectra were recorded with 256 scans and are plotted after dividing by the noise rmsd and $\sqrt{ns}$ to facilitate a direct comparison. Apodization is not used.

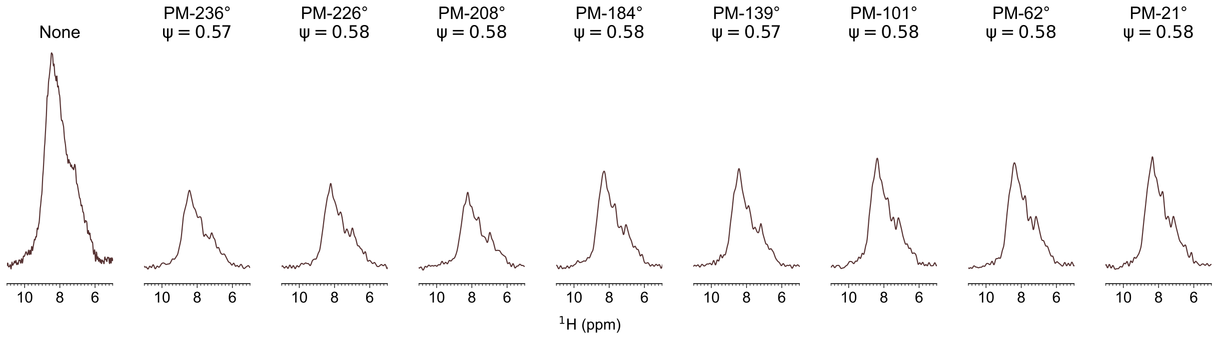


FIG. S23. $^{15}$N edited $^{1}$H 1D spectra of GB1 recorded at 60 kHz MAS on a 1.3 mm HCN probe without homonuclear decoupling (No windowed detection) and with homonuclear decoupling with windowed detection with phase ramps 236°($\psi$ = 0.57, rf = 125 kHz, $\lambda$ = 0.81, NS = 32), 226°($\psi$ = 0.58, rf = 125 kHz, $\lambda$ = 0.82, NS = 32), 208°($\psi$ = 0.58, rf = 120 kHz, $\lambda$ = 0.79, NS = 32), 184°($\psi$ = 0.57, rf = 90 kHz, $\lambda$ = 0.8, NS = 32), 139°($\psi$ = 0.57, rf = 55 kHz, $\lambda$ = 0.83, NS = 32), 101°($\psi$ = 0.58, rf = 45 kHz, $\lambda$ = 0.85, NS = 32), 62°($\psi$ = 0.58, rf = 40 kHz, $\lambda$ = 0.84, NS = 32), 21°($\psi$ = 0.58, rf = 40 kHz, $\lambda$ = 0.82, NS = 32). The spectrum without homonuclear decoupling was recorded with 8 scans using standard acquisition. The spectra with homonuclear decoupling are scaled by a chemical shift scaling factor ($\lambda$). Spectra are plotted after dividing by the noise rmsd and $\sqrt{ns}$ to facilitate a direct comparison.

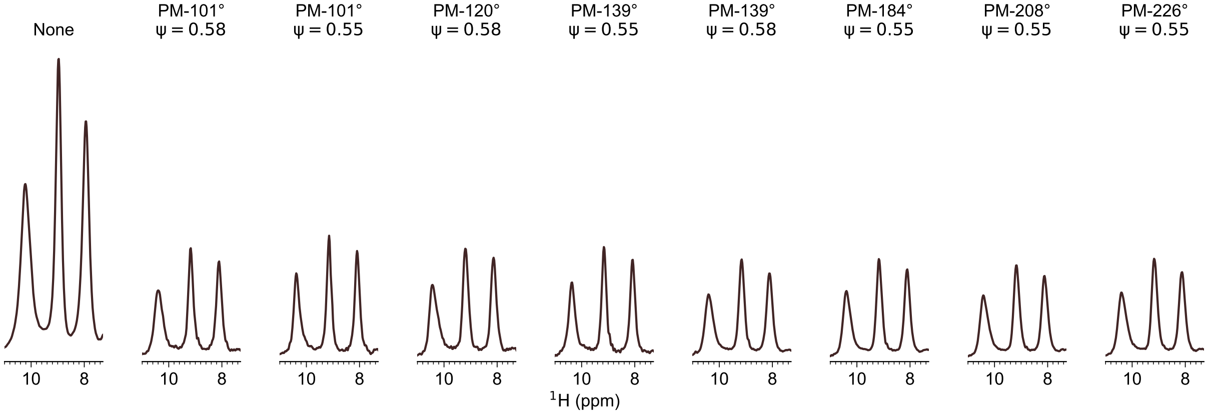


FIG. S24. $^{15}$N edited $^{1}$H 1D spectra of f-MLF recorded at 95.2 kHz MAS on 0.7 mm HCN probe to compare the sensitivity of without homonuclear decoupling (No windowed detection) and with homonuclear decoupling with windowed detection with phase ramps 101°($\psi$ = 0.58, rf = 175 kHz, $\lambda$ = 0.84, NS = 128), 101°($\psi$ = 0.55, rf = 109 kHz, $\lambda$ = 0.89, NS = 128), 120°($\psi$ = 0.55, rf = 117 kHz, $\lambda$ = 0.89, NS = 1024), 120°($\psi$ = 0.58, rf = 206 kHz, $\lambda$ = 0.84, NS = 128), 139°($\psi$ = 0.55, rf = 149 kHz, $\lambda$ = 0.86, NS = 128), 139°($\psi$ = 0.58, rf = 170 kHz, $\lambda$ = 0.89, NS = 1024), 184°($\psi$ = 0.55, rf = 185 kHz, $\lambda$ = 0.89, NS = 1024), 208°($\psi$ = 0.55, rf = 219 kHz, $\lambda$ = 0.89, NS = 1024), and 226°($\psi$ = 0.55, rf = 219 kHz, $\lambda$ = 0.92, NS = 1024). The spectrum without homonuclear decoupling was recorded with 1024 scans(NS). All the spectra with homonuclear decoupling are plotted after scaling by the chemical shift scaling factor($\lambda$). Spectra are plotted after dividing by the noise rmsd and $\sqrt{ns}$ to facilitate a direct comparison. Apodization is not used.

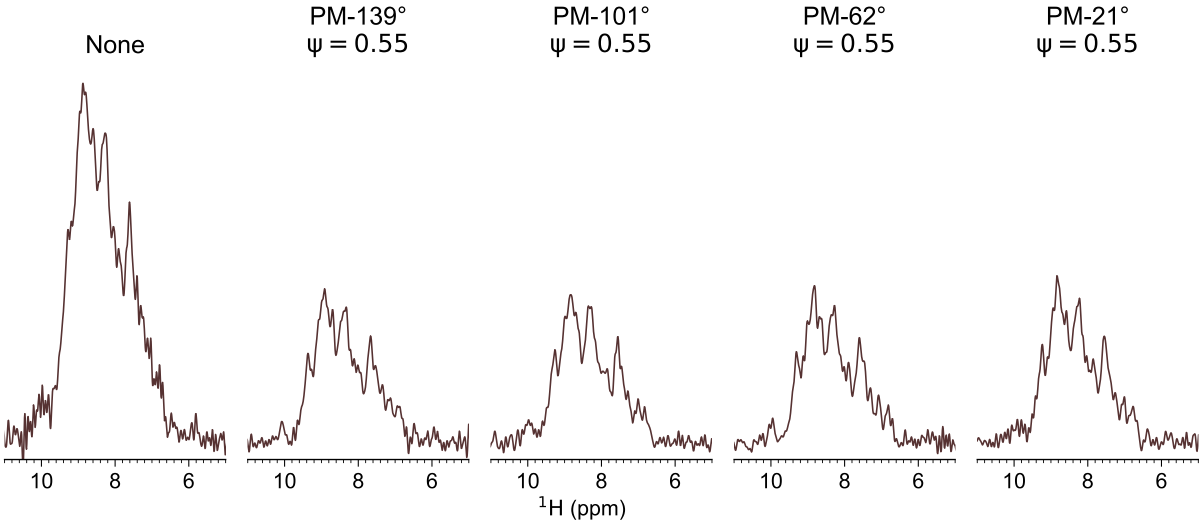


FIG. S25. $^{15}$N edited $^{1}$H 1D spectra of GB1 recorded at 95.24 kHz MAS on a 0.7 mm HCN probe without homonuclear decoupling (No windowed detection) and with homonuclear decoupling with windowed detection with phase ramps 139°($\psi$ = 0.55, rf = 140 kHz, $\lambda$ = 0.87, NS = 32), 101°($\psi$ = 0.55, rf = 110 kHz, $\lambda$ = 0.83, NS = 32), 62°($\psi$ = 0.55, rf = 100 kHz, $\lambda$ = 0.88, NS = 32), and 21°($\psi$ = 0.55, rf = 90 kHz, $\lambda$ = 0.89, NS = 32). The spectrum without homonuclear decoupling was recorded with 8 scans. The spectra with homonuclear decoupling are scaled by a chemical shift scaling factor ($\lambda$). Spectra are plotted after dividing by the noise rmsd and $\sqrt{ns}$ to facilitate a direct comparison.

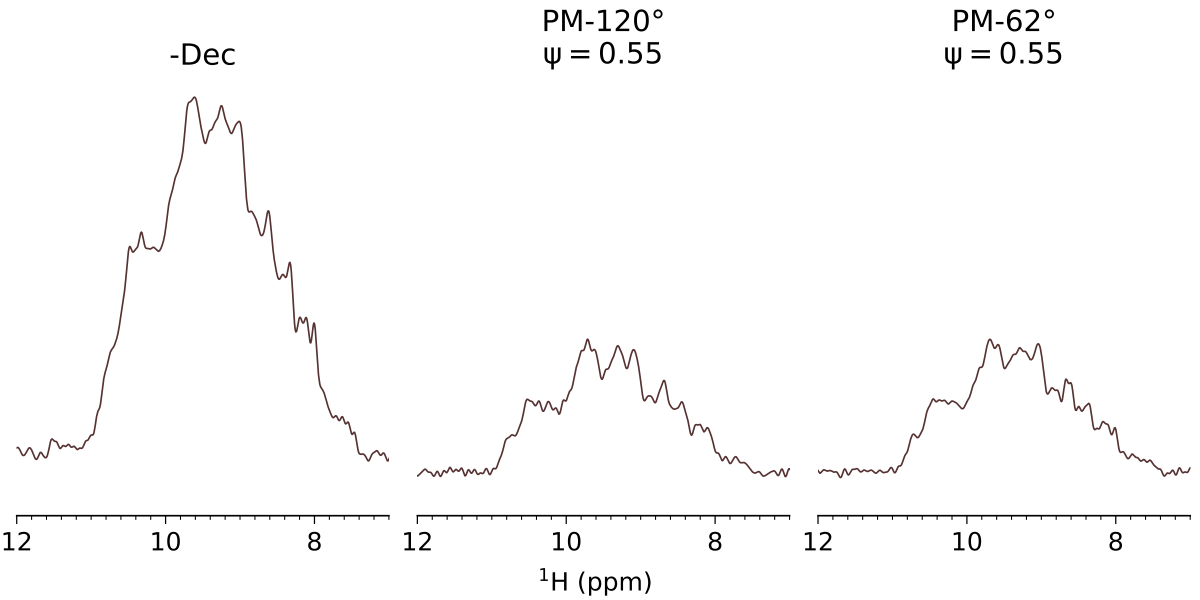


FIG. S26. $^{15}$N edited $^{1}$H 1D spectra of ParM recorded at 95.24 kHz MAS on a 0.7 mm HCN probe without homonuclear decoupling (No windowed detection) and with homonuclear decoupling with windowed detection with phase ramps 120°($\psi$ = 0.55, rf = 124 kHz, $\lambda$ = 0.9, NS = 128) and 62°($\psi$ = 0.55, rf = 82 kHz, $\lambda$ = 0.932, NS = 128). The spectrum without homonuclear decoupling was recorded with 32 scans. The spectra with homonuclear decoupling are scaled by a chemical shift scaling factor($\lambda$). Spectra are plotted after dividing by the noise rmsd and $\sqrt{ns}$ to facilitate a direct comparison.

### S7. PULSE SEQUENCES FOR BRUKER SPECTROMETERS

#### A. $^{15}$N-edited $^{1}$H for optimising homonuclear decoupling

```
; #---channels
# define H f1
# define C f2
# define N f3

; #---default delays
  "d58 = 1m"
  "d59 = 200u"
  "d60 = 0.1u"
  "d61 = 0.5u"
  "d62 = 1.0u"
  "d63 = 10u"

; #---misc
# ifdef xdec
  "acqt0 = d61 + d60"
# else
  "acqt0 = 0u"
# endif

; #---mas
  define delay taur
  "taur = 1s/cnst50"

; #---decoupling
# ifdef rcwhp
;p32: 180 deg pulse for rcw (high power) decoupling
;p31: CW pulse for rCW-ApA decoupling (set internally)
  "p31 = (0.98*taur) - (0.5*p32)"
# endif

# ifdef rcwlp
;l12: N (14/16/18)*taur
;p32: pulse for low power rCW-ApA decoupling (set internally)
;p31: pulse for low power rCW-ApA decoupling (set internally)
  "p31 = 0.2125 * l12 * taur"
  "p32 = 0.075 * l12 * taur"
# endif

; #---homonuclear decoupling
;p18: tau_LG
;l19: number of pulses in pmlg (10 for m5m, 6 for m3m
;p19: length of a single pulse
;d3: window duration
;spnam18: homonuclear decoupling (m5m, m3m, times)

;cnst18: omega_c / omega_r
  "p18 = l19 * p19"

; #---d18 is a constant echo
;d18: echo time
;l18: loops for echo (set by d18)
# ifdef nowindow
  "l18 = d18 / (4 * p18)"
```

```
  "cnst18 = 1e6 / (2 * p18 * cnst50)"
# else
  "l18 = d18 / (4 * (p18 + d3))"
  "cnst18 = 1e6 / (2 * (p18 + d3) * cnst50)"
# endif

; #--------------------------------------------------#

; This  section of this pp is variable
; check the stored pp for the actual values used

; #---2d optimization for power and pulse length
;l6: length of the the power array
; calibrated on MLF 0-100 kHz in steps of 10 kHz
  define list<power> pwl = {Watt 0.000 0.057 0.230 0.523 0.937 1.462 2.128 2.893 3.755 4.776 5.945}
  define loopcounter ni
  "ni = td1 / l6"

; #--------------------------------------------------#

Check, d63
  ze
  d18
  "d8 = cnst18*1u"
  d8

  d63 pwl.res

Start, d63

; #---set new homdec power and pulse length
  "p18 = l19 * p19"

; #---recalculate number of loops to maintain constant echo time
# ifdef nowindow
  "l18 = d18 / (4 * p18)"
# else
  "l18 = d18 / (4 * (p18 + d3))"
# endif

; #---recycle delay
  d1
  d63 fq=0:N
  d63 pl1:H

; #---90 pulse
  (p1 ph1):H

; #---CP
  (p13:sp13 ph20):H (p13:sp34 ph15):N
  (p3 pl3 ph21):N

; #---water suppression
  d60 pl17:H pl3:N
  d61 fq=cnst17:H
  d60 cpds7:H
  d17
  d61 fq=0:H
```

```
; #---bring X back
  (p3 ph11):N
  d60 do:H

; #---reverse cp
  (p31:sp16 ph20):H (p31:sp31 ph18):N

; #---homodec+echo
  d60 pwl:H

# ifdef nowindow
3 (p18:sp18(currentpower) ph10^):H
  (p18:sp18(currentpower) ph10^):H
  lo to 3 times l18
  d60

  (p1*2 pl1 ph2):H

  d60 pwl:H
4 (p18:sp18(currentpower) ph10^):H
  (p18:sp18(currentpower) ph10^):H
  lo to 4 times l18

# else

3 d3*0.5
  (p18:sp18(currentpower) ph10^):H
  d3
  (p18:sp18(currentpower) ph10^):H
  d3*0.5
  lo to 3 times l18
  d60

  (p1*2 pl1 ph2):H

  d60 pwl:H
4 d3*0.5
  (p18:sp18(currentpower) ph10^):H
  d3
  (p18:sp18(currentpower) ph10^):H
  d3*0.5
  lo to 4 times l18

# endif

; #--detect
  d60 pl33:N
  go=Start ph31 cpds3:N finally do:N

  d58 wr #0 if #0 zd

  d63 pwl.inc
  lo to Start times l6

  d63 pwl.res
  d63 ipu19
  lo to Start times ni
```

```
HaltAcqu, d58
exit

; #---phase tables
; min=16, 16*n, maybe 8 if you are stretched

  ph1  = 1
  ph15 = 0
  ph11 = 3 1
  ph18 = 0 0 2 2
  ph10 = 0 2
  ph2  = 0 0 0 0 1 1 1 1 2 2 2 2 3 3 3 3
  ph31 = 0 2 2 0 2 0 0 2

  ph20 = 0
  ph21 = 1
  ph22 = 2
  ph23 = 3
; #--------------------
```

#### B. $^{15}$N-$^{1}$H HSQC using windowed detection

```
; hNH_win
; hNH correlation with C decoupling during NH evolution and windowed detection with homonuclear decoupling

# define H f1
# define C f2
# define N f3

; #---mas
  define delay taur
  "taur = 1s / cnst50"

; #---defult delays
  "d63 = 10u"
  "d62 = 2u"
  "d61 = 0.5u"
  "d60 = 0.1u"
  "d58 = 1.0m"
  "acqt0 = d60"

; #---windowed pmlg settings
#include <Avancesolids.incl>
#include <Delayssolids.incl>
  "d9 = 0.1u * (l11)"       ; set the sampling window in Avancesolids.incl
  define delay window
  "window = p9"              ; defines the widow in conjuction with d9
  "p18 = l19 * p19"          ; wPMLG pulse length
  "d4 = window - (p1 / 2)"

  define delay dead
  "dead = window - d9 - d8 - 0.1u"   ; part of the window

  define delay cycle
  "cycle = 2 * (window + p18)"

  define loopcounter count
  "count = aq / cycle"  ;make sure td datapoints are sampled
  "blktr2 = 0.7u"       ;this opens the transmitter gate 0.7 usec before the
                        ;pulse, so the transmitter noise is not sample
  "anavpt = l11"        ;for analog mode

  define delay dwell_1H
  "dwell_1H = cycle / 2"
  "cnst6 = 1/(dwell_1H)"
  "d6 = cnst6 * 1u"
  "cnst9 = 1/ (cnst50 * cycle)"
  "l31 = (aq / dw)"

;cnst9: PSI = (omega_c / omega_r)
;cnst6: Spectral width that showld be put in SWH [Hz] in the direct dimension
;count: make sure TD ~ 4*count

# ifdef rcwhp
;p52: 180 deg pulse for rcw (high power) decoupling
;p51: CW pulse for rCW-ApA decoupling (set internally)
  "p51 = (0.98*taur) - (0.5*p52)"
```

```
# endif

# ifdef rcwlp
;l12: N =14/16/18
;p52: pulse for low power rCW-ApA decoupling (set internally)
;p51: pulse for low power rCW-ApA decoupling (set internally)
  "p51 = 0.2125 * l12 * taur"
  "p52 = 0.075 * l12 * taur"
# endif

; #---2d statements
  "in0 = inf1"
# ifdef oned
  "d0 = 0.1u"
# else
  "d0 = (in0 / 2) - (d60 * 4)"
# endif
# ifdef cnst_duty
  "d30 = d0 + (td1 - 1) * in0 / 2 + 10u"
# endif

; #---------#
; #  START  #
; #---------#

Pre, ze

  d63 fq=cnst6:H
  d63 fq=cnst9:H
  d63 fq=0:H

Check, d63

# ifndef dont_check_sw
   if "l31 > ((count * 4) - 16)" goto passTD1
  print "SWH is probably set incorrectly to a value lower than expected. It should be equal to cnst6"
  goto HaltAcqu
passTD1, d60

   if "l31 < ((count * 4) + 2)" goto passTD2
  print "SWH is probably set incorrectly to a value higher than expected. It should be equal to cnst6"
  goto HaltAcqu
passTD2, d60
# endif

  if "taur < 100u" goto pass_mas
  print "taur too long (>100u). Is MAS < 10kHz?"
  goto HaltAcqu
pass_mas, d60

  if "p13 < 10m" goto pass_p13
  print "fowrward cp (p13) too long (>10m). Aborting"
  goto HaltAcqu
pass_p13, d60

  if "p31 < 10m" goto pass_p31
  print "reverse cp (p31) too long (>10m). Aborting"
  goto HaltAcqu
```

```
pass_p31, d60

  if "aq < 30m" goto pass_aq
  print "aq too long (>30m). Aborting"
  goto HaltAcqu
pass_aq, d60

  if "d17 < 200m" goto pass_d17
  print "d17 too long (>200m). Aborting"
  goto HaltAcqu
pass_d17, d60

# ifndef oned
  if "d0 + ((td1 - 2) * in0 / 2) < 50m" goto pass_d0
  print "final d0 too long (>50m). Aborting"
  goto HaltAcqu
pass_d0, d60
# endif

# ifndef oned
  if "aq + d17 + d0 + (((td1 / 2) - 1) * in0) < (0.3 * d1)" goto pass_duty
  print "duty cycle too long (> 30%). Aborting"
  goto HaltAcqu
pass_duty, d60
# endif

Start, d1

; #---for homonuclear decoupling
  d63 reset1:H      ;synchronise pulse and detection RF
  STARTADC          ;prepare adc for sampling, set reference frequency, defined in Avancedru.incl
  RESETPHASE        ;reset reference phase (ph30)
  d63 rpp10         ;reset phase list pointer

; #---90 cp
  (p1 pl1 ph1):H
# ifdef cnst_duty
  (p13:sp13 ph20):H (p13:sp34 ph16 p3 pl3 ph21):N
# else
  (p13:sp13 ph20):H (p13:sp34 ph16):N
# endif

# ifndef oned
; #---15N evolution
  d60 pl11:H
  d60 cpds1:H
# ifdef cnst_duty
  d30
  (p3 pl3 ph23):N
  d60
  d60
# endif
  (center (d0) (p2 pl2 ph20 p2*2 pl2 ph21 p2 pl2 ph20):C)
  d60
  d60
# endif
  (p3 pl3 ph3):N
  d60 do:H
```

```
; #---water suppression
  d62 fq=cnst17:H
  d60 pl17:H
  d60 cpds7:H
  d17
  d60 do:H
  d62 fq=0:H

; #---reverse cp
# ifdef calN90
  (p39 pl39 ph4):N
# else
  (p3 pl3 ph4):N
# endif
  (p31:sp16 ph20):H (p31:sp31 ph5):N

; #---detect
  d60 pl33:N
  d60 cpds3:N

; #---windowed acquisition starts
4 dead
  d8 RG_ON
  sample
  (p18:sp18 ph10^):H
  dead
  d8 RG_ON
  sample
  (p18:sp18 ph10^):H
  lo to 4 times count
; #---windowed acquisition ends
  d60 do:N

; #---loop for ns scans with phase table increments
  d63 rcyc=Start

; #---2D
  d58 mc #0 to Start
  F1PH(calph(ph3, +90), caldel(d0, +in0) & caldel(d30, -in0))


HaltAcqu, d63
exit

; #---phase tables
;ns: 2, 4, 8, 16, 32, 16*n, 32*n
  ph1  = 1 1 1 1 3 3 3 3
  ph16 = {0}*16 {2}*16
  ph3  = 1 1 3 3
  ph4  = {3}*8 {1}*8
  ph5  = 0 2
  ph10 = 0 2
  ph30 = 0
  ph31 = 0 2 2 0 2 0 0 2
         2 0 0 2 0 2 2 0
         2 0 0 2 0 2 2 0
         0 2 2 0 2 0 0 2
```

```
; #---default phases
  ph20 = 0
  ph21 = 1
  ph22 = 2
  ph23 = 3
```

#### C. Shape File for Homonuclear Decoupling

A representative shape file for homonuclear decoupling (PM-120$^o$) is provided below:

```
##TITLE=PM_120
##JCAMP-DX= 5.00 Bruker JCAMP library
##DATA TYPE= Shape Data
##ORIGIN= Bruker BioSpin GmbH
##OWNER= <BRUKER>
##DATE= 2005/11/29
##TIME= 14:47:39
##$SHAPE_PARAMETERS=
##MINX= 1.000000E02
##MAXX= 1.000000E02
##MINY= 2.560000E01
##MAXY= 3.256000E02
##$SHAPE_EXMODE= None
##$SHAPE_TOTROT= 0.000000E00
##$SHAPE_TYPE= Excitation
##$SHAPE_USER_DEF=
##$SHAPE_REPHFAC=
##$SHAPE_BWFAC= 0.000000E00
##$SHAPE_BWFAC50=
##$SHAPE_INTEGFAC= 6.534954E-17
##$SHAPE_MODE= 0
##NPOINTS= 6
##XYPOINTS= (XY..XY)
1.000000E02 325.60
1.000000E02 265.60
1.000000E02 205.60
1.000000E02 25.60
1.000000E02 85.60
1.000000E02 145.60
##END
```

[1] Franks, W. T. *et al.* Magic-angle spinning solid-state NMR spectroscopy of the beta1 immunoglobulin binding domain of protein G (GB1): $^{15}$N and $^{13}$C chemical shift assignments and conformational analysis. *J. Am. Chem. Soc.* **127**, 12291–12305 (2005).

[2] Taware, P. P. *et al.* Measuring dipolar order parameters in nondeuterated proteins using solid-state NMR at the magic-angle-spinning frequency of 100 kHz. *J. Phys. Chem. Lett.* **14**, 3627–3635 (2023).

[3] Raran-Kurussi, S., Sarawata, B., Balasubramaniam, D. & Mote, K. R. A comparison between MBP and NT* as N-terminal fusion partner for recombinant protein production in E. coli. *Protein Expr. Purif.* **189**, 105991 (2022).
